# Knocking out the carboxyltransferase interactor 1 (CTI1) in Chlamydomonas boosted oil content by fivefold without affecting cell growth

**DOI:** 10.1101/2024.09.03.611075

**Authors:** Zhongze Li, Minjae Kim, Jose Roberto da Silva Nascimento, Bertrand Legeret, Gabriel Lemes Jorge, Marie Bertrand, Fred Beisson, Jay J Thelen, Yonghua Li-Beisson

## Abstract

The first step in chloroplast *de novo* fatty acid synthesis is catalyzed by acetyl-CoA carboxylase (ACCase). As the rate-limiting step for this pathway, ACCase is subject to both positive and negative regulation. In this study, we identify a Chlamydomonas homolog of the plant carboxyltransferase interactor 1 (CrCTI1) and show that this protein, interacts with the Chlamydomonas α-carboxyltransferase (Crα-CT) subunit of the ACCase by yeast two-hybrid protein-protein interaction assay. Three independent CRISPR-Cas9 mediated knock-out mutants for CrCTI1 each produced an “enhanced oil” phenotype, accumulating 25% more total fatty acids and storing up to five-fold more triacylglycerols (TAGs) in lipid droplets. The TAG phenotype of the *crcti1* mutants was not influenced by light but was affected by trophic growth conditions. By growing cells under heterotrophic conditions, we observed a crucial function of CrCTI1 in balancing lipid accumulation and cell growth. Mutating a previously mapped *in vivo* phosphorylation site (CrCTI1 Ser108 to either Ala or to Asp), did not affect the interaction with Crα-CT. However, mutating all six predicted phosphorylation sites within Crα-CT to create a phosphomimetic mutant reduced significantly this pairwise interaction. Comparative proteomic analyses of the *crcti1* mutants and WT suggested a role for CrCTI1 in regulating carbon flux by coordinating carbon metabolism, antioxidant and fatty acid β-oxidation pathways, to enable cells adapt to carbon availability. Taken together, this study identifies CrCTI1 as a negative regulator of fatty acid synthesis in algae and provides a new molecular brick for genetic engineering of microalgae for biotechnology purposes.

## INTRODUCTION

Microalgae, thanks to their capacity to convert CO_2_ into reduced carbon and stored in the form of starch or oils, are considered a sustainable feedstock for lipid production (Hu *et al*., 2008). However, simultaneously satisfying both cell growth and lipid accumulation is a great challenge since the amount of lipid content and growth rate are often inversely correlated (Williams and Laurens, 2010). Understanding the regulation of lipid biosynthetic mechanisms may help formulate additional strategies to improve lipid productivity in microalgae.

As the building blocks of glycerolipids, fatty acids are produced by the *de novo* fatty acid synthesis in the chloroplasts (Chapman and Ohlrogge, 2012). The initial step, ATP-dependent carboxylation of acetyl-CoA to produce malonyl-CoA is catalyzed by acetyl-CoA carboxylase (ACCase). In green algae and most higher plants including dicots and nongraminaceous monocots, the chloroplastic ACCase is a heteromeric complex that contains four catalytic subunits named biotin carboxylase, biotin carboxyl carrier protein, and α- and β-carboxyltransferases (CTs) (Guchhait *et al*., 1974; Konishi and Sasaki, 1994). As the rate-limiting step in *de novo* fatty acid synthesis, ACCase is highly regulated through different mechanisms; such as stroma pH, ATP/ADP ratio, Mg^2+^, redox potential, and feedback regulation by downstream metabolites like 18:1-acyl carrier protein (Davis and Cronan, 2001; Kleinig and Liedvogel, 1979; Roesler *et al*., 1997; Sauer and Heise, 1983). In addition, several proteins are reported as negative regulators for ACCase activity, including PII and BADC proteins (Gerhardt *et al*., 2015; Salie *et al*., 2016; Zalutskaya *et al*., 2015).

In a study in *Arabidopsis thaliana*, proteins called Carboxyltransferase Interactors (CTIs) were discovered as negative regulators of ACCase in plant leaves (Ye *et al*., 2020). These small proteins, located in the chloroplast inner envelope membrane, were found to regulate ACCase activity by interacting with the α-CT subunit through the coiled-coil domain at its C-terminus (Ye *et al*., 2020). The authors suggested that the association of ACCase with CTI may be light-dependent based on the phenotypes of CTI knockout mutants including characterization of protein interaction, activity status and pathway flux, as well as accumulation of higher TAGs and total fatty acids under light adaptation (Ye *et al*., 2020).

To test whether a similar negative regulator of fatty acid synthesis occurs in microalgae, and to pinpoint the molecular mechanisms underlying the regulation of ACCase by CTI, we identified the occurrence of a single CTI homolog in the unicellular green microalga *Chlamydomonas reinhardtii* (hereafter Chlamydomonas). Furthermore, we report the isolation of knock-out mutants (named *crcti1*) of the CTI homolog using CRISPR-Cas9 method. The CTI homolog (CrCTI1) present in Chlamydomonas is confirmed to interact with the α-CT subunit of ACCase through yeast two-hybrid (Y2H) assay. Investigation of the Chlamydomonas *crcti1* mutants under different cultivation conditions (i.e., photoauto-, mixo- and heterotrophy) demonstrated that CrCTI1 functions as a negative regulator of *de novo* fatty acid synthesis in green microalgae, and that control of CrCTI1 on ACCase is not directly dependent on light, in contrast to the plant ortholog. Moreover, we show that the absence of CrCTI1 during optimal mixotrophic growth improves TAG content by at least 5-fold without compromising cell growth.

## RESULTS

### Identification of only one CTI in Chlamydomonas

In Arabidopsis, three CTI homologs have been identified and characterized: AtCTI1 (At1g42960), AtCTI2 (At3g02900), and AtCTI3 (At5g16660) (Ye *et al*., 2020). Using BLASTp analysis, we found only one CTI homolog (Cre06.g278195) in the Chlamydomonas and named it CrCTI1 (**Figure 1A**). This protein has previously been named Chloroplast Punctuate Protein 1 (CPP1) since it has been localized to chloroplast and appeared as punctate distribution (Wang *et al*., 2023). To avoid confusion, hereafter, we used the name referring their function; CrCTI1. Domain analysis for secondary structure indicated that CrCTI1 protein contains a putative N terminal chloroplast transit peptide (1–35 AA), a single transmembrane domain (43– 66 AA) followed by a coiled-coil domain (93–118 AA) at the C-terminal region (**Figure 1B**). Overall, the domain structure was similar to AtCTI1, but the length of CrCTI1 was shorter than that of AtCTI1 due to both shorter chloroplast transit peptide and coiled-coil domains. The protein-protein interactions between CrCTI1 and Crα-CT were predicted using the alpha-fold program, indicating that they likely interact at their respective coiled-coil domains (**Figure 1C**).

**Figure 1.**
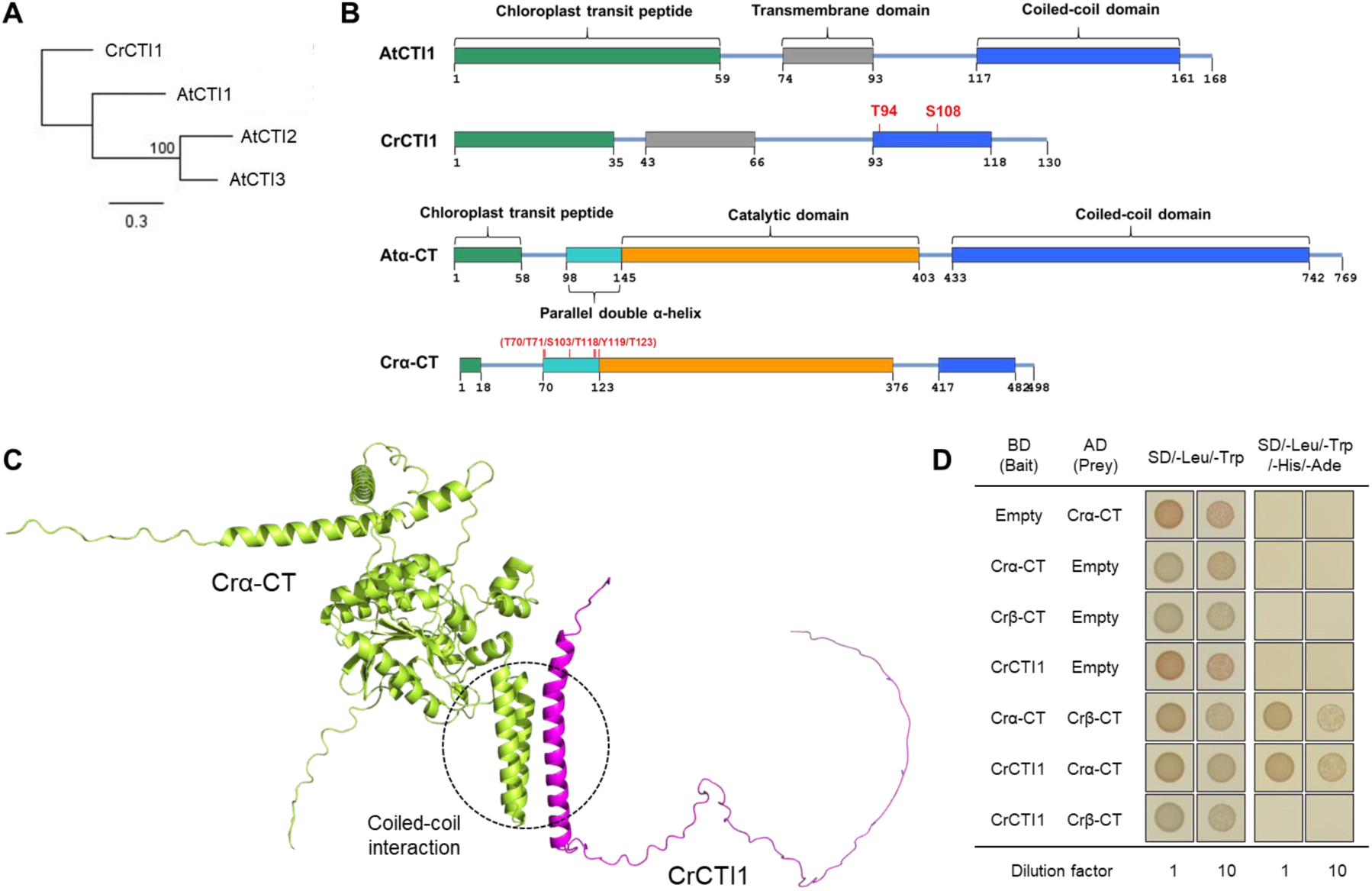
Chlamydomonas contains one ortholog of the Arabidopsis CTIs which interacts with the α-CT subunit of ACCase. **(A)** Phylogenetic tree of CrCTI1 and three Arabidopsis CTI homologs constructed by comparing amino acid sequences. **(B)** Protein domain structures of *A. thaliana* CTI1 (AtCTI1), *C. reinhardtii* CTI1 (CrCTI1), *A. thaliana* α-CT (Atα-CT), and *C. reinhardtii* α-CT (Crα-CT). The location of putative phosphorylation sites was highlighted in red color. **(C)** Prediction model of coiled-coil interaction (highlighted by dot circle) between Crα-CT (light green) and CrCTI1 (magenta) using AlphaFold2. **(D)** Yeast two-hybrid (Y2H) assay showed that full-length Crα-CT interacts with the coiled-coil domain of CrCTI1 *in vitro*. Empty vector and Crβ-CT served as negative controls, the test between Crα-CT and Crβ-CT served as positive control. AD, pGADT7 (prey vector); BD, pGBKT7 (bait vector); SD, synthetic drop-out medium supplements.

### CrCTI1 interacts with the α-CT subunit of the Chlamydomonas ACCase

The previous study in Arabidopsis discovered that AtCTIs specifically interact with the α-CT subunit of ACCase (Ye *et al*., 2020). To investigate the possible interaction of a putative CrCTI1 with Chlamydomonas ACCase, we first investigated protein-protein interactions with both the α- and β-CT subunits from Chlamydomonas. For the Y2H assay, the coiled-coil domain-containing C-terminal region (68-130 AA) of the CrCTI1 was cloned into a bait vector (pGBKT7; BD) and full length Crα-CT or Crβ-CT were cloned into a prey vector (pGADT7; AD). Protein-protein interaction was validated based on cell viability in different nutritional culture plates: synthetic drop-out (SD) medium lacking leucine and tryptophan (SD/-Leu/-Trp), or SD medium lacking leucine, tryptophan, histidine, and adenine (SD/-Leu/-Trp/-His/-Ade). Colony formation should be observed on the plate of SD/-Leu/-Trp/-His/-Ade when there is an interaction between the proteins in AD and BD. There was no colony formation in the combination of CrCTI1, Crα-CT, and Crβ-CT with an empty vector (**Figure 1D**). As a component of the ACCase complex, α-CT is known to bind to β-CT, so they were used as a positive control. As expected, we could observe the colony formation from the combination of Crα-CT and Crβ-CT (**Figure 1D**). Among several combinations of AD and BD tested, CrCTI1 interacted only with Crα-CT (**Figure 1D**).

### Phosphorylation state of the CrCTI1 does not seem to affect its interaction with Crα-CT

Next, we attempted to investigate what are the regulatory factors dictate the interaction between CrCTI1 and Crα-CT. Protein phosphorylation is one well-known regulatory mechanism for protein-protein interaction by causing either charge or conformational changes which can affect binding strength (Nishi *et al*., 2011). In Arabidopsis, mining of multiple phospho-proteomic databases showed that AtCTI1 and Atα-CT are both phosphorylated (Agrawal and Thelen, 2006; Meyer *et al*., 2012; Umezawa *et al*., 2013). Based on this, we hypothesized that phosphorylation could potentially play a role in the regulation of this interaction. Chlamydomonas in vivo phosphoproteomic data showed phosphorylation of Serine 108 (S108) within the coiled-coil domain of CrCTI1 (CrCTI1_CC) (**Supplemental Figure S1A**)(Roustan *et al*., 2017). To investigate whether phosphorylation of the S108 residue could affect its interaction with Crα-CT, we generated the yeast strains expressing phosphomimetic mutation (S108D) or non-phosphomimetic mutation (S108A) of the CrCTI1-CC domain. The Y2H assay on plate and liquid culture showed that phosphorylation state of the S108 does not affect the interaction (**Supplemental Figures S1B** and **S1C**).

To further expand the repertoire of phosphorylation-mediated interaction, we searched whether it occurs other phosphorylation targets. Using the phosphorylation site prediction tool NetPhos-3.1 (Blom *et al*., 1999), additional putative phosphorylation sites were identified as CrCTI1_CC (T94) and Crα-CT (T70, T71, S103, T118, Y119, and T123) (**Figure S1A**). We mutated all putative phosphorylation sites either all to Asp (D) or all to Ala (A). Along with wild type protein (native), these mutated versions were transformed into yeast. Interestingly, the combination with phosphomimetic mutation of Crα-CT (Crα-CT (D)) indicated a significantly lower number of colony formation compared to others (**Figure S1D**). These results suggest that the phosphorylation state of the Crα-CT may influence the interaction with CrCTI1.

### Generation and validation of CrCTI1 knockout mutants using CRISPR-Cas9

To study the function of CrCTI1 in microalgae, we generated knockout mutants in the Chlamydomonas CC4349 strain (wild type, hereafter WT) using the CRISPR-Cas9 RNP method. The single guide RNA (sgRNA) was designed to target the 2^nd^ exon of the *CrCTI1* gene (**Figure 2A**). The RNP complex and the hygromycin-resistance (*HygR*) gene cassette were co-transformed into cells. Colonies grown on agar plates containing hygromycin were screened for mutants by genomic DNA PCR against the sgRNA target sequence (**Figure 2B**). The insertion of the *HygR* gene was validated by Sanger sequencing (**Figure 2C**). RT-PCR analysis showed that there is no transcription of *CrCTI1* in the mutants (**Figure 2D**). We thus named the three independent transformants as *crcti1-1*, *crcti1-2*, and *crcti1-3*.

**Figure 2.**
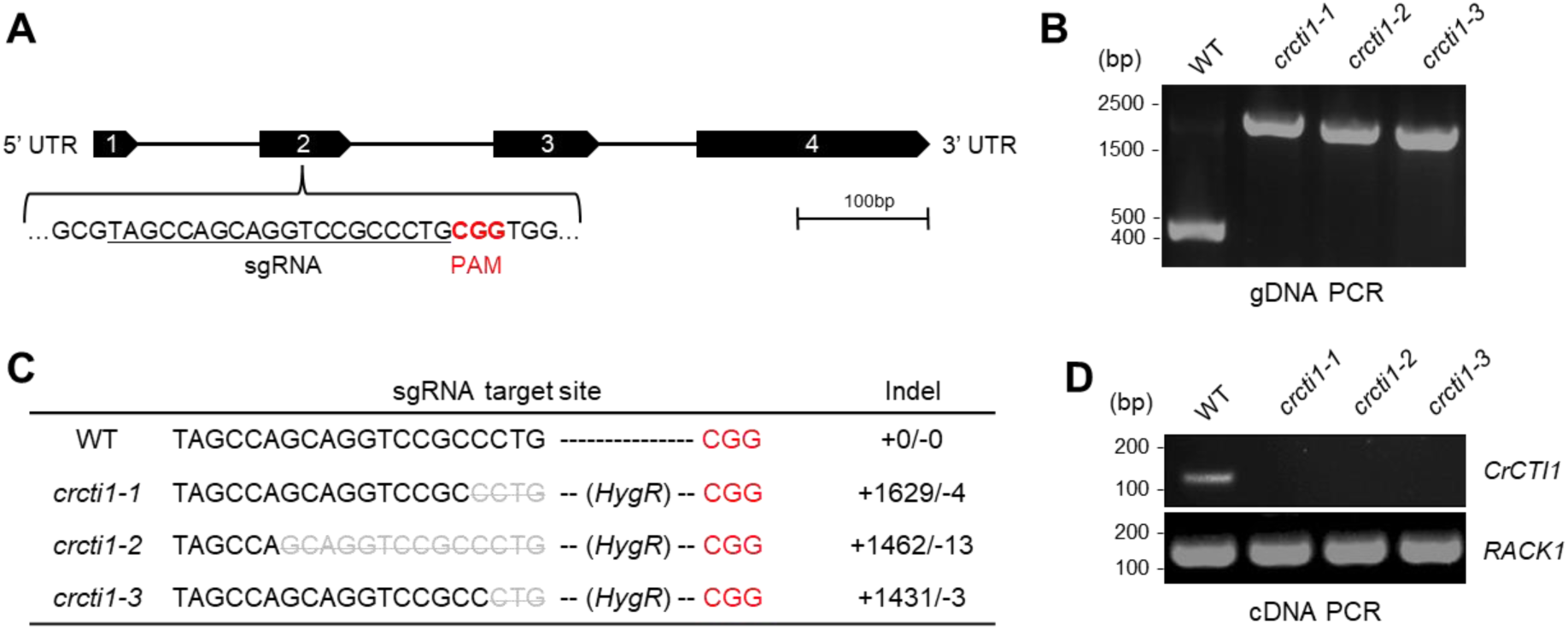
Generation and validation of knock-out mutants for Chlamydomonas CrCTI1 using CRISPR-Cas9. **(A)** The sgRNA target site on *CrCTI1* gene. Black arrow boxes with number stand for exons, and the black line represent the introns. **(B)** Genomic DNA (gDNA) PCR verification of cassette insertion in the *crcti1* mutants. **(C)** Sanger sequencing results. **(D)** RT-PCR analyses of *CrCTI1* expression in WT and the three *crcti1* mutants. *RACK1*, receptor for activated kinase C 1 was used as reference gene. M, marker; WT, wild type; PAM: protospacer adjacent motif.

### Chlamydomonas *crcti1* mutants make 5-fold more oil than WT during mixotrophic growth

To investigate the role of CrCTI1 in lipid synthesis, we measured the lipid amount and fatty acid composition of the cells grown under mixotrophic conditions (light, ambient CO**_2_**, and acetate). The total fatty acid content of *crcti1* mutants was more than 25% higher than WT; *crcti1-1* (1.55-fold), *crcti1-2* (1.25-fold), and *crcti1-3* (1.57-fold) (**Figure 3A**). The desaturation ratio was decreased in the *crcti1* mutants (**Supplemental Figure S2A**). Despite the relative low TAG content (< 1 μg per 10^6^ cells) of Chlamydomonas cells during optimal growth, the *crcti1* mutants exhibited significantly higher TAG content than WT: *crcti1-1* (7.11-fold), *crcti1-2* (4.65-fold), and *crcti1-3* (6.96-fold) (**Figure 3B**). Consistently, confocal microscopic images indicated that *crcti1* mutant cells have a clear lipid droplet accumulation compared to WT (**Figure 3C**). TAGs can be synthesized through *de novo* TAG synthesis as well as membrane lipid remodeling (Young and Shachar-Hill, 2021). Thus, we measured polar membrane lipid classes to assess whether membrane lipid turnover is different between WT and the mutants. Thin layer chromatography (TLC) analysis of polar lipids showed that they are all at similar levels between WT and the three *crcti1* mutants (**Supplemental Figure S2B**), suggesting that improved TAG amount in the mutants is mostly likely due to increased *de novo* fatty acid synthesis and unlikely due to enhanced membrane lipid turnover. But of course, we are measuring steady state level of lipid amount, and thus can not rule out the possibility of more rapid lipid turnover in the *crcti1* mutants.

**Figure 3.**
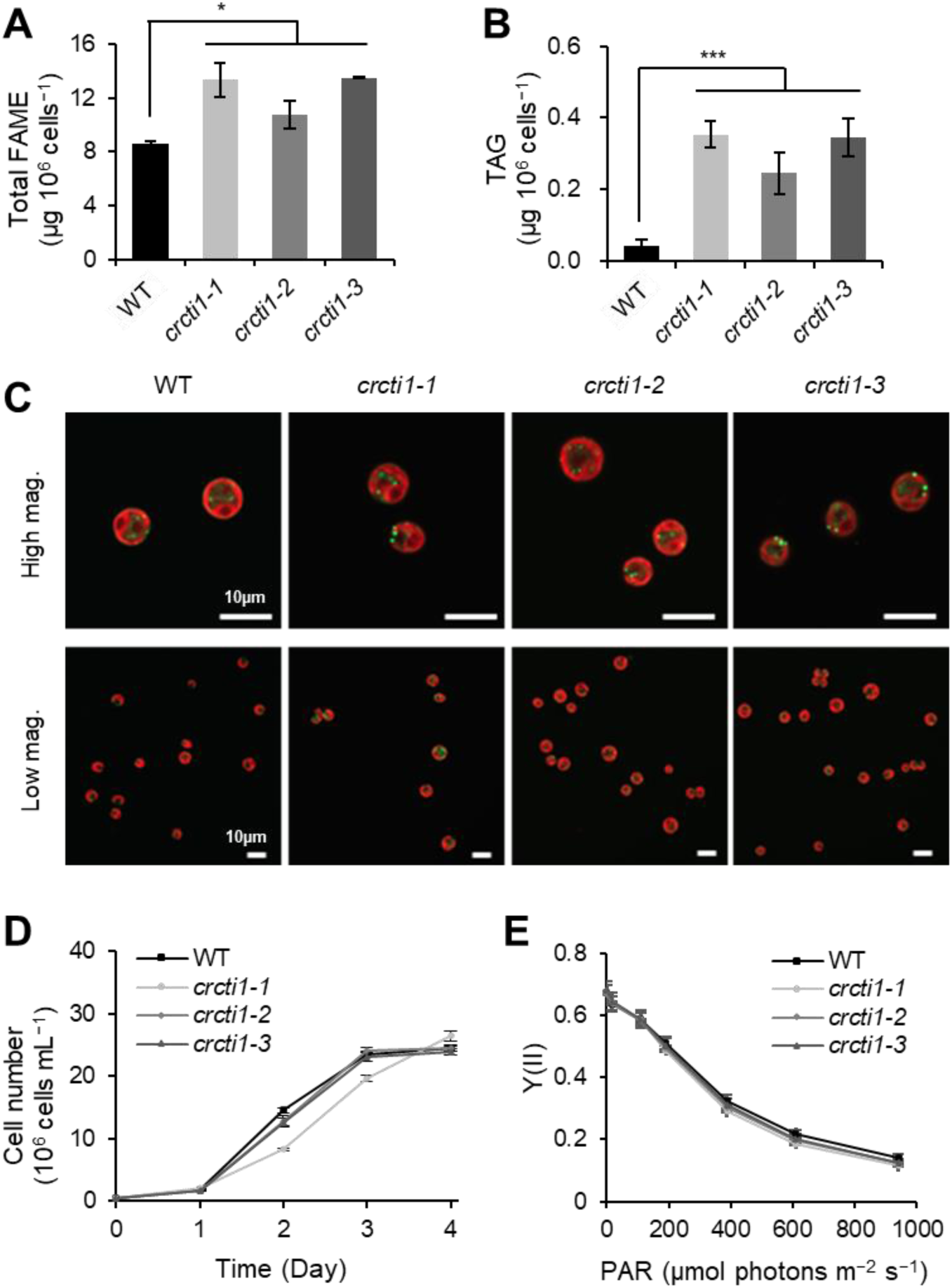
Chlamydomonas CrCTI1 mutants made more oil during mixotrophic growth without compromising photosynthesis yield and growth. **(A)** Total fatty acids content measured by GC-MS. FAME, fatty acid methyl ester. **(B)** TAG content measured by thin layer chromatography (TLC). **(C)** Confocal fluorescence microscopy images of Chlamydomonas cells. Green, Bodipy fluorescence; red, chlorophyll autofluorescence. Images were present in high and low magnification. Scale bars = 10 µm. **(D)** Chlamydomonas cell growth curve. **(E)** PSII yield (Y(II)) measurement. Data are means of three biological replicates with standard deviation shown. \**p* < 0.05, \*\*\**p* < 0.001 (Student’s *t*-test).

In addition, cell growth was similar between all strains (**Figure 3D**), and this is consistent with photosynthetic activity as measured by chlorophyll fluorescence (**Figure 3E**). Therefore, *crcti1* mutants made significantly higher amounts of TAGs without compromising growth during mixotrophic growth on acetate. Since nitrogen (N) starvation is known to enhance *de novo* fatty acid synthesis and TAG accumulation, we also analyzed fatty acid and TAG amounts of the three *crcti1* mutants under nitrogen starved conditions. We found that the *crcti1* mutants accumulated at least 40% more TAGs than WT (**Supplemental Figure S2C**), suggesting that CrCTI1 exerts negative control on *de novo* fatty acid synthesis regardless of nitrogen status. Taken together, knocking out the CrCTI1 in Chlamydomonas has shown an additive effect to further boost TAG accumulation under N starvation.

### Oil accumulation in the *crcti1* mutants is affected by CO_2_ level in the environment

To further define the physiological conditions when CrCTI1 exerts its function, we analyzed cell growth and lipid amount under photoautotrophic conditions when CO_2_ is the sole carbon source for growth.

First, there was no difference in photosystem II efficiency under both conditions (**Supplemental Figure S3**). As we observed in the mixotrophic condition, the total fatty acid content of *crcti1* mutants was higher than WT regardless of the CO_2_ level in the air (**Figure 4A**). The altered fatty acid composition showed a significant decrease in the relative proportion of 16:4, 18:3, and 18:4 polyunsaturated fatty acids and an increase of 16:2 and 18:2. These results indicate a reduced desaturation ratio in all the *crcti1* mutant lines under photoautotrophic conditions (**Supplemental Figure S4**), consistent with the idea that *de novo* fatty acid synthesis is enhanced. Interestingly, there was no significant difference in TAG content under high CO_2_ conditions, whereas the *crcti1* mutants accumulated 2.5-fold more TAG than WT under ambient CO_2_ conditions (**Figure 4B**). This is not surprising considering the high growth rate observed under high versus low CO_2_ (air), i.e. under high CO_2_, a higher demand for fatty acids to meet the requirement of membrane synthesis in WT (**Figure 4C** versus **Figure 4D**). There is indeed an overall higher amount of total fatty acids per cell under high CO_2_ than air (**Figure 4A**). The *crcti1* mutants and WT showed similar growth behavior under high CO_2_ conditions, but the mutants grew slower than WT under ambient CO_2_ conditions (**Figure 4C**). These results indicate that CrCTI1 could play a role in regulating cell growth and storage lipid synthesis under limited CO_2_ availability.

**Figure 4.**
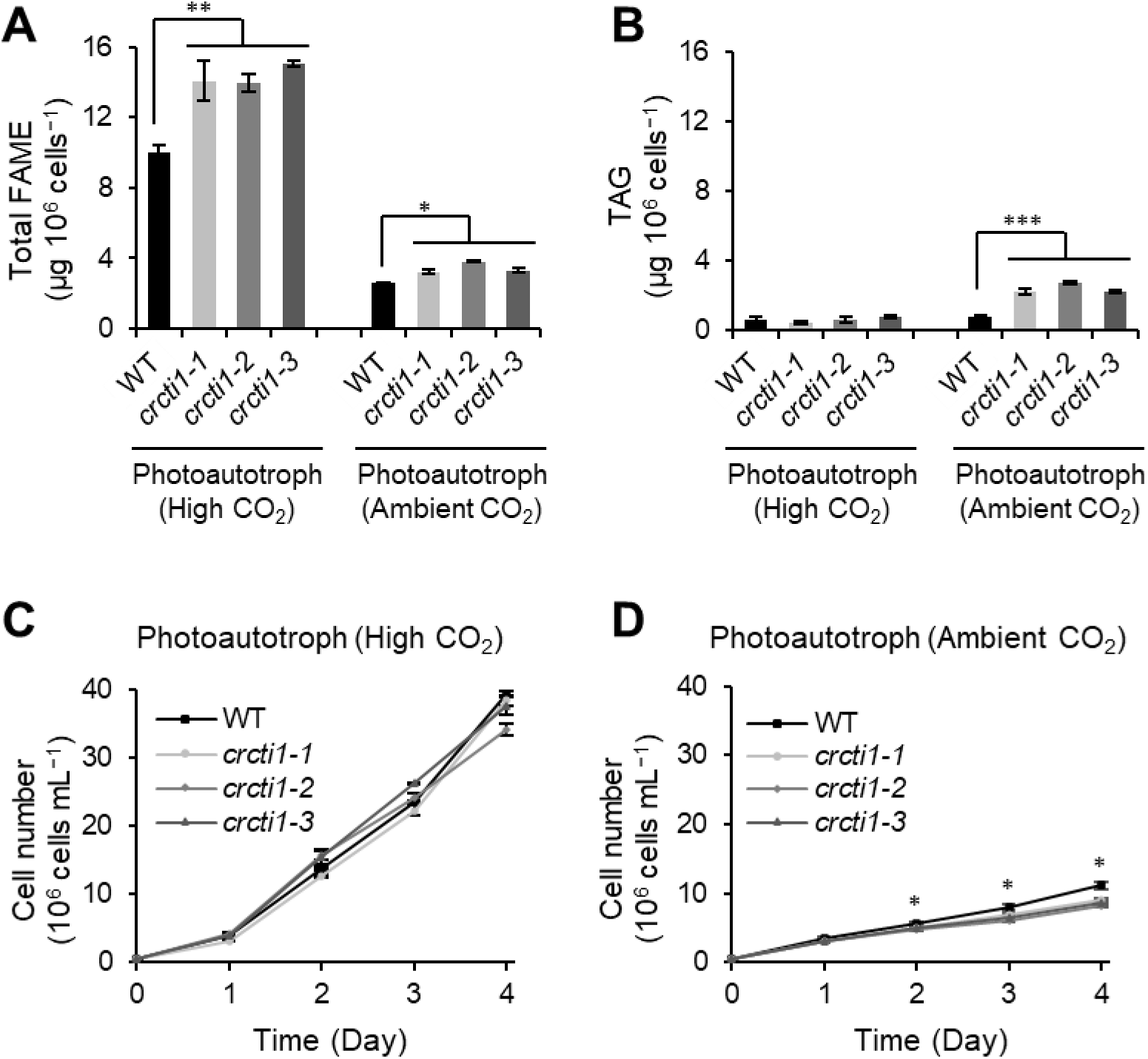
Oil accumulation in *cti1* mutants is affected by CO_2_ level during photoautotrophic growth. **(A)** Total fatty acids content measured by GC-MS. **(B)** TAG content measured by thin layer chromatography (TLC). **(C)** Cell growth curve cultured in photoautotrophic condition with additional 2% CO_2_ supply in the air. **(D)** Cell growth curve cultured in photoautotrophic condition with ambient air. All experiments were performed in three biological replicates (± S.D). * *p* < 0.05, ** *p* < 0.01, *** *p* < 0.001 (Student’s *t*-test). MM, minimal medium. FAME, fatty acid methyl ester.

### Physiological characterization of *crcti1* mutants under heterotrophic conditions

In Arabidopsis, it was reported that the negative regulation of fatty acid synthesis by CTI in the leaves is light-dependent (Ye *et al*., 2020). As a photoautotrophic organism, Arabidopsis fix CO_2_ as the sole carbon source for growth. Unlike Arabidopsis, the green microalga Chlamydomonas can grow under photoautotrophic, mixotrophic as well as heterotrophic conditions when the cells can directly utilize external organic carbon like acetate as the only carbon source in the dark. We thus posed the question whether CrCTI1 had a role in the regulation of fatty acid synthesis in Chlamydomonas in the dark. To answer this question, we first cultivated all 4 strains in mixotrophic conditions until the mid-log phase, then these cells were sub-cultured and kept in the dark with acetate for three days before cells were harvested for lipid analysis (**Figure 5A**).

**Figure 5.**
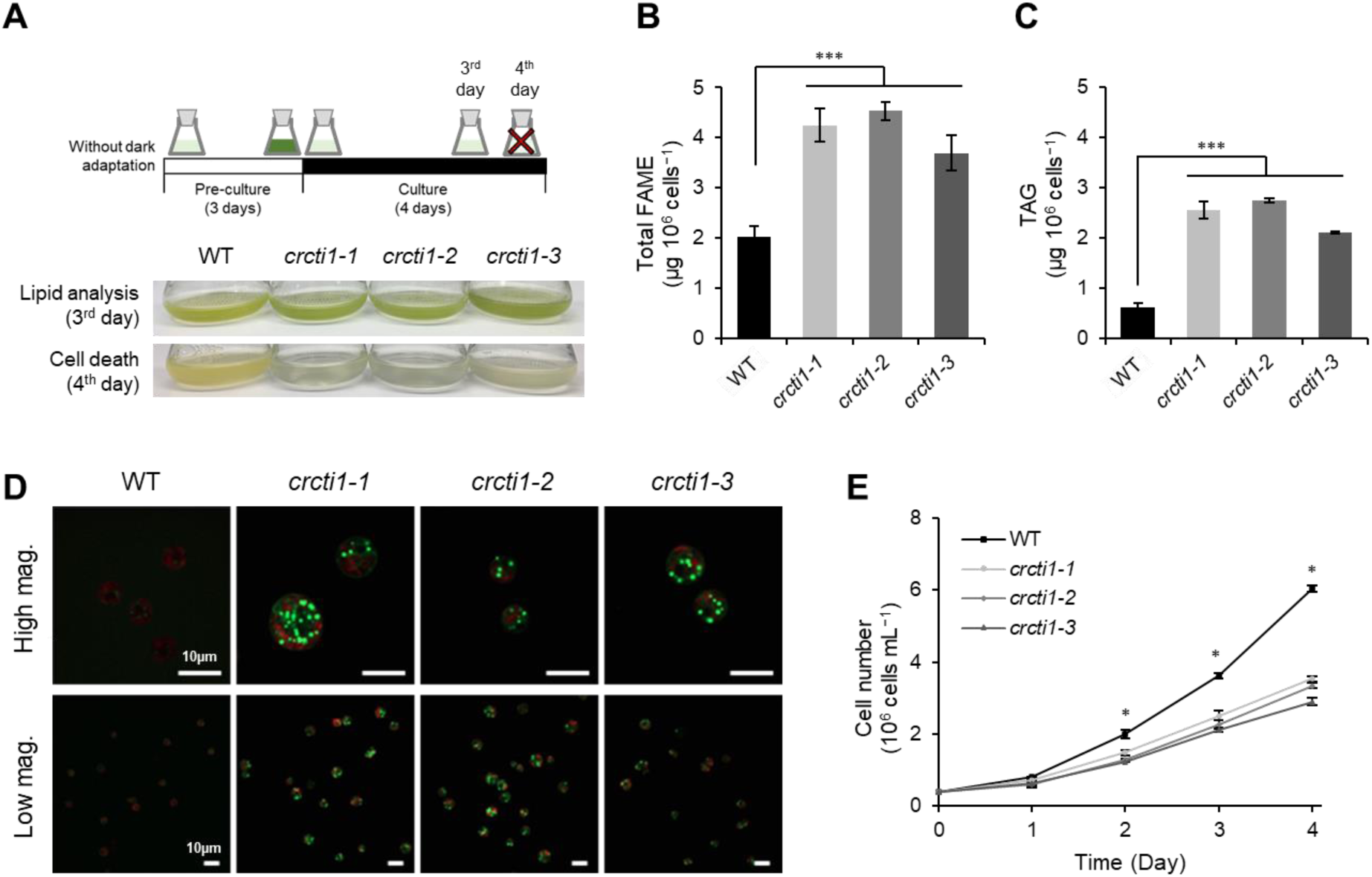
Physiological characteristics of *crcti1* mutants during heterotrophic growth. **(A)** Working flow of the experiments and the pictures of cell culture in flasks. Images were taken on the 3 d of pre-culture or at the 4th d of sub-culture in the dark. **(B)** Total fatty acids content measured by GC-MS. FAME, fatty acid methyl ester. **(C)** TAG content measured by thin layer chromatography (TLC). **(D)** Confocal fluorescence microscopy images of Chlamydomonas cells. Green, Bodipy fluorescence; red, chlorophyll autofluorescence. Images were present in high and low magnification. Scale bars = 10 µm. **(E)** Chlamydomonas cell growth curve cultured in TAP medium in the dark. All experiments were performed in three biological replicates (± S.D). * *p* < 0.05, *** *p* < 0.001 (Student’s *t*-test).

During heterotrophic growth, we observed that the total fatty acid content of the *crcti1* mutants was almost twice higher than WT; *crcti1-1* (2.09-fold), *crcti1-2* (2.22-fold), and *crcti1-3* (1.82-fold) (**Figure 5B**). In addition, TAG content of the *crcti1* mutants was around 4-fold higher than WT; *crcti1-1* (4.13-fold), *crcti1-2* (4.44-fold), and *crcti1-3* (3.40-fold) (**Figure 5C**). Confocal microscopy images showed a clear increase in both the number and size of lipid droplets in *crcti1* mutants comparing to WT (**Figure 5D**). In addition, we observed that the growth of *crcti1* mutants was significantly repressed under heterotrophic condition (**Figure 5E**). Even more dramatic, after being cultivated in TAP in the dark for an extended period (after 4 days), the sub-culture of mutant cells could not grow and lost viability (**Figure 5A**).

### Global proteomics reveals metabolic remodeling as a result of *crcti1* mutation

Global comparative proteomics has proven useful for elucidating changes in metabolic pathways between plants with differing oil-content genotypes (Chen and Pramanik, 2009; Hajduch *et al*., 2007). Here we analyzed the protein abundance in the *crcti1* mutants under mixotrophic and photoautotrophic conditions (air level or enriched with 2% CO_2_); the heterotrophic condition was excluded because cell densities were too low to obtain sufficient amount of proteins. The proteome dataset was produced from four biological replicates for each of the three *crcti1* mutants and WT (**Supplemental Table S1**).

To be considered a differential protein it must be observed in at least two *crcti1* mutant alleles (Log2(*crcti1*/WT) ≥ 1 or Log2(*crcti1*/WT) ≤ -1) (**Supplemental Figure S5**). In this analysis, protein LHCP (refer to nuclear-encoded light-harvesting chlorophyll- and carotenoid-binding protein), translocation defect (LTD, Cre12.g551950 (Jeong *et al*., 2018)), and monodehydroascorbate reductase 1 (MDAR1, Cre17.g712100) were identified as up-regulated proteins in *crcti1* mutants under all three conditions (**Supplemental Figure S5A**). Phosphoenolpyruvate carboxylase 1 (Cre16.g673852) was down-regulated in *crcti1* mutants under all three conditions (**Supplemental Figure S5B**). A more comprehensive list of differential proteins is presented in **Supplemental Figures 6A, 6B, and 6C** showing the 50 most variable proteins in each condition (TAP, MM air, MM CO_2_, respectively).

**Figure 6** shows average of log2FC for the three mutant alleles (*crcti1*/WT) for proteins involved in central carbon metabolism. The data show that *crcti1* mutations impact various aspects of central carbon metabolism in a manner influenced by available acetate. Acetyl-CoA synthase (ACS) converts acetate into acetyl-CoA, playing a crucial role in acetate assimilation. Chlamydomonas harbors three ACS isoforms, ACS1 (Cre01.g071662), ACS2 (Cre01.g055408, orthologous to Arabidopsis chloroplast ACS), and ACS3 (Cre07.g353450), which has been experimentally demonstrated to be localized in peroxisomes (Lauersen *et al*., 2016). Chloroplast ACS2 was more abundant under mixotrophic conditions, while peroxisomal ACS3 was reduced under both mixotrophic and photoautotrophic (ambient CO_2_) conditions and increased under photoautotrophic (high CO_2_) growth conditions. Regarding peroxisomal fatty acid catabolism, the peroxisomal fatty acid transporter PXA1 (Cre15.g637761), peroxisomal acyl-CoA synthase LCS3 (Cre12.g507400) (Bai *et al*., 2021), and enoyl-CoA hydratase 1 ECH3 (Cre16.g695050) were each reduced under mixotrophic growth. These data suggest that fatty acid degradation is repressed under mixotrophic conditions and partly explains the higher fold increase in TAG accumulation under mixotrophic growth in the *crcti1* mutants (**Figure 3**).

**Figure 6.**
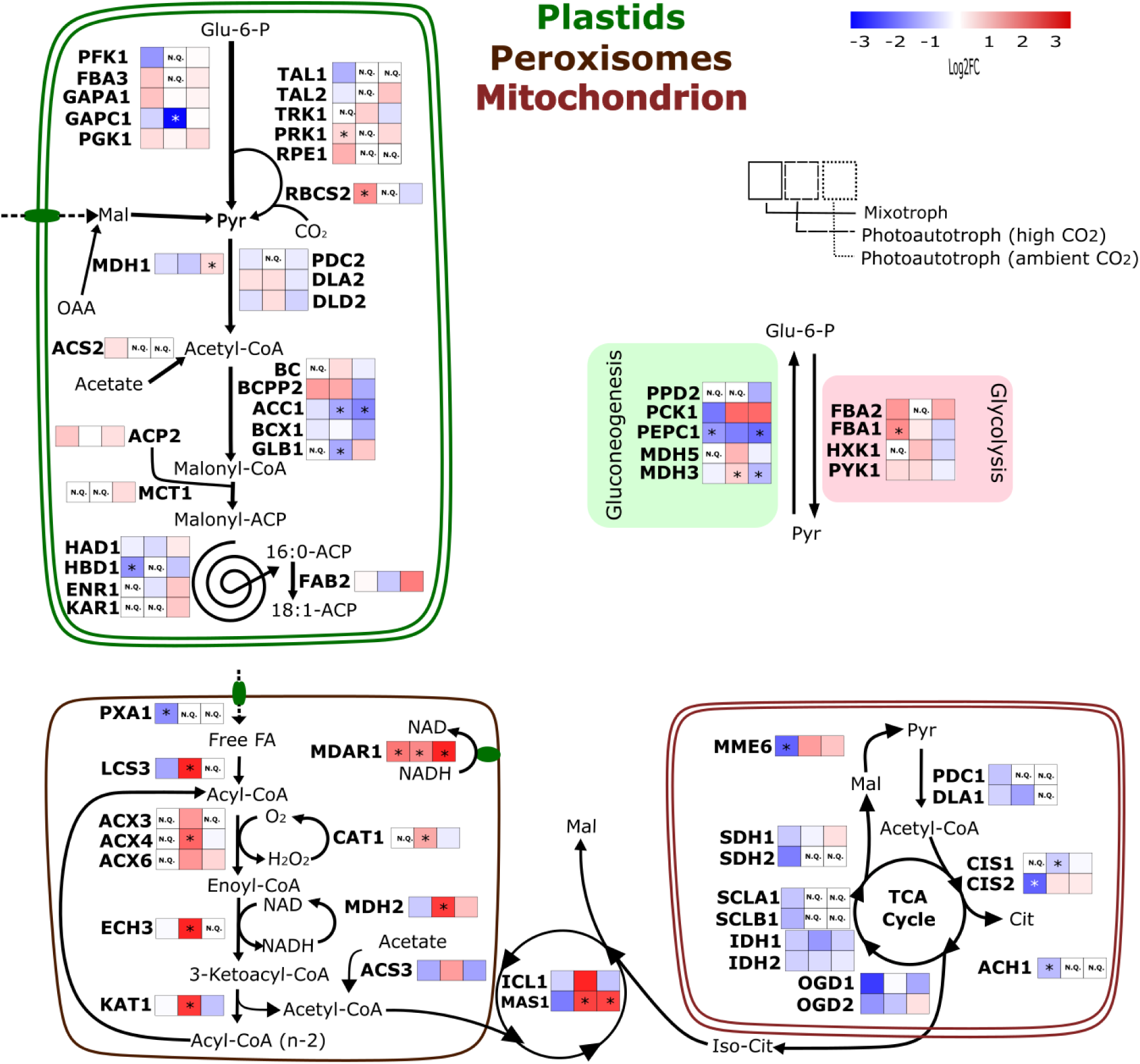
Metabolic adaptations as a result of *crcrti1* mutation under mixotrophic and photoautotrophic conditions. Fold-changes of protein amounts between CrCTI1 mutants and wild-type (average log2FC between all three mutants – *crcti1*/WT) grouped according to pathways. N.Q. represent proteins not differently abundance with a *p* value ≤ 0.05. * represent two to more mutant lines with *p*-value ≤ 0.01, and similar fold-change values. Abbreviations: ACC1, alpha-carboxyltransferase (ACCase complex); ACH1, aconitate hydratase; ACLB1, ATP citrate lyase, subunit B; ACP2, acyl-carrier protein 2; ACS2, acetyl-CoA synthase 2; ACS3, acetyl-CoA synthetase/ligase 3; ACX3, acyl-CoA oxidase/dehydrogenase 3; ACX4, putative acyl-CoA oxidase 4; ACX6, acyl-CoA oxidase/dehydrogenase 6; BC1, biotin carboxylase (ACCase complex); BCC1, acetyl-CoA biotin carboxyl carrier; BCX1, beta-carboxyltransferase (ACCase complex); CAT1, mono-functional catalase; CIS1, citrate synthase, mitochondrial; CIS2, citrate synthase, glyoxysomal/microbody form; DLA1, dihydrolipoamide acetyltransferase; DLA2, dihydrolipoamide acetyltransferase; DLD2, dihydrolipoamide dehydrogenase 2; ECH3, enoyl-CoA hydratase 1; ENR1, enoyl-ACP reductase; FAB2, plastid acyl-ACP desaturase; FBA1, fructose-1,6-bisphosphate aldolase; FBA2, fructose-1,6-bisphosphate aldolase; FBA3, fructose-1,6-bisphosphate aldolase, chloroplastic; FRK2, probable fructokinase-2; GAPA1, glyceraldehyde-3-phosphate dehydrogenase A; GAPC1, glyceraldehyde 3-phosphate dehydrogenase; GAPC, chloroplastic; GLB1, nitrogen regulatory protein PII; HAD1, 3-hydroxyacyl-ACP dehydratase; HBD1, 3-hydroxybutyrate dehydrogenase; HXK1, hexokinase; IDH1, isocitrate dehydrogenase, NAD-dependent; IDH2, isocitrate dehydrogenase, NAD-dependent; KAR1, 3-oxoacy-ACP reductase; KAT1, 3-oxoacyl CoA thiolase/acetyl-CoA acyltransferase 1; LCS3, long-chain acyl-CoA synthetase; MAS1, malate synthase; MCT1, malonyl-CoA:acyl-carrier-protein transacylase; MDAR1, monodehydroascorbate reductase 1; MDH1, NAD-dependent malate dehydrogenase 1, chloroplastic; MDH2, malate dehydrogenase 2; MDH3, NAD-dependent malate dehydrogenase 3; MDH5, NADP-malate Dehydrogenase 5; MME6, NADP-dependent malic enzyme 6; OGD1, 2-oxoglutarate dehydrogenase, E1 subunit; OGD2, dihydrolipoamide succinyltransferase, oxoglutarate dehydrogenase E2 component; PCK1, phosphoenolpyruvate carboxykinase; PDC1, mitochondrial pyruvate dehydrogenase complex, E1 component, alpha subunit; PFK1, phosphofructokinase; PGK1, phosphoglycerate kinase, chloroplastic; PPD2, pyruvate phosphate dikinase; PRK1, phosphoribulokinase, chloroplastic; PYK1, pyruvate kinase 1; RBCS2, ribulose-1,5-bisphosphate carboxylase/oxygenase (RuBisCO) small subunit 2, chloroplastic; RPE1, ribulose phosphate-3-epimerase, chloroplastic; SBE3, starch branching enzyme 3; SCLA1, succinyl-CoA ligase alpha chain; SCLB1, succinate-CoA ligase beta chain; SDH1, succinate dehydrogenase flavoprotein subunit; SDH2, iron-sulfur subunit of mitochondrial succinate dehydrogenase; TAL1, transaldolase; TAL2, putative transaldolase, plastid form; TRK1, transketolase, chloroplastic.

β-oxidation was differentially induced under photoautotrophic conditions depending on CO_2_ levels. All β-oxidation enzymes increased under high CO_2_ but not under ambient CO_2_ which showed no change. This agrees with the contrasting fold change levels in fatty acids and TAG measured for photoautotrophic high and ambient CO_2_ (**Figure 4A, B**). Peroxisomal acetyl-CoA, produced by β-oxidation, is incorporated into organic acids by the glyoxylate cycle, and the abundance of the two enzymes (ACS2 and the peroxisomal malate synthase1 MAS1 (Cre03.g144807) (Lauersen *et al*., 2016) and isocitrate lyase ICL1 (Cre06.g282800), exclusively involved in the glyoxylate cycle further supports fatty acid degradation influence on TAG levels in *crcti1* mutants under mixotrophic and photoautotrophic high CO_2_ conditions, especially as MAS1, and ICL1 were among the twenty-five most upregulated proteins under photoautotrophic (high CO_2_) conditions.

Chloroplast carbon metabolism was also altered in *crcti1* mutants. HetACCase subunits α-CT and β-CT both decreased under all conditions, though more intensely under photoautotrophic conditions. Chloroplast glycolysis phosphoglycerate kinase 1 PGK1 (Cre11.g467770), fructose-bisphosphate aldolase 3 FBA3 (Cre05.g234550), and glyceraldehyde-3-phosphate dehydrogenase GAPA1 (Cre01.g010900) mostly increased under both mixotrophic and photoautotrophic conditions. The reductive pentose phosphate pathway was also impacted; RuBisCO RBCS2 (Cre02.g120150), phosphoribulokinase PRK1 (Cre12.g554800), and ribulose-5-phosphate-3-epimerase RPE1 (Cre12.g511900) all increased under mixotrophic conditions.

Mitochondrial carbon metabolism was also differently altered in *crcti1* mutants under different conditions. All quantified TCA cycle enzymes and mitochondrial malic enzyme were reduced under mixotrophic conditions. These data suggest the *crcti1* mutant is shifting carbon from TCA intermediate production to fatty acid synthesis. Considering TCA cycle provides carbon precursors for amino acid and carbohydrate biosynthesis, this putative metabolic shift agrees with the higher increase in TAG in *crcti1* mutants under mixotrophic conditions.

### Increase in enzymatic antioxidant defense pathway can effectively compensate for oxidative stress

FA β-oxidation produces H_2_O_2_ through the activity of acyl-CoA oxidases (Kong *et al*., 2017). To compare the oxidative state of WT and *crcti1* mutants, we measured the DCFH-DA fluorescence as an indicator of intracellular H_2_O_2_ levels (Yoshida *et al*., 2003). Interestingly, there was no difference in H_2_O_2_ levels under mixotrophic and photoautotrophic (high CO_2_) conditions where sufficient carbon was available, whereas the *crcti1* mutants were found to have higher H_2_O_2_ levels than WT under photoautotrophic conditions (ambient CO_2_) (**Figure 7A**). Next, we investigated the protein abundance for the enzymes related to the intracellular antioxidant defense pathway (**Figure 7B**). MDAR1 regenerates ascorbate in the enzymatic antioxidant defense pathway of plants (Eltayeb *et al*., 2007). Because MDAR1 was more abundant in *crcti1* mutants under both mixotrophic and photoautotrophic conditions, we hypothesized that the lack of CrCTI1 places the mutant under higher oxidative pressure and that the increase in MDAR1 is a response to this increased oxidative pressure.

**Figure 7.**
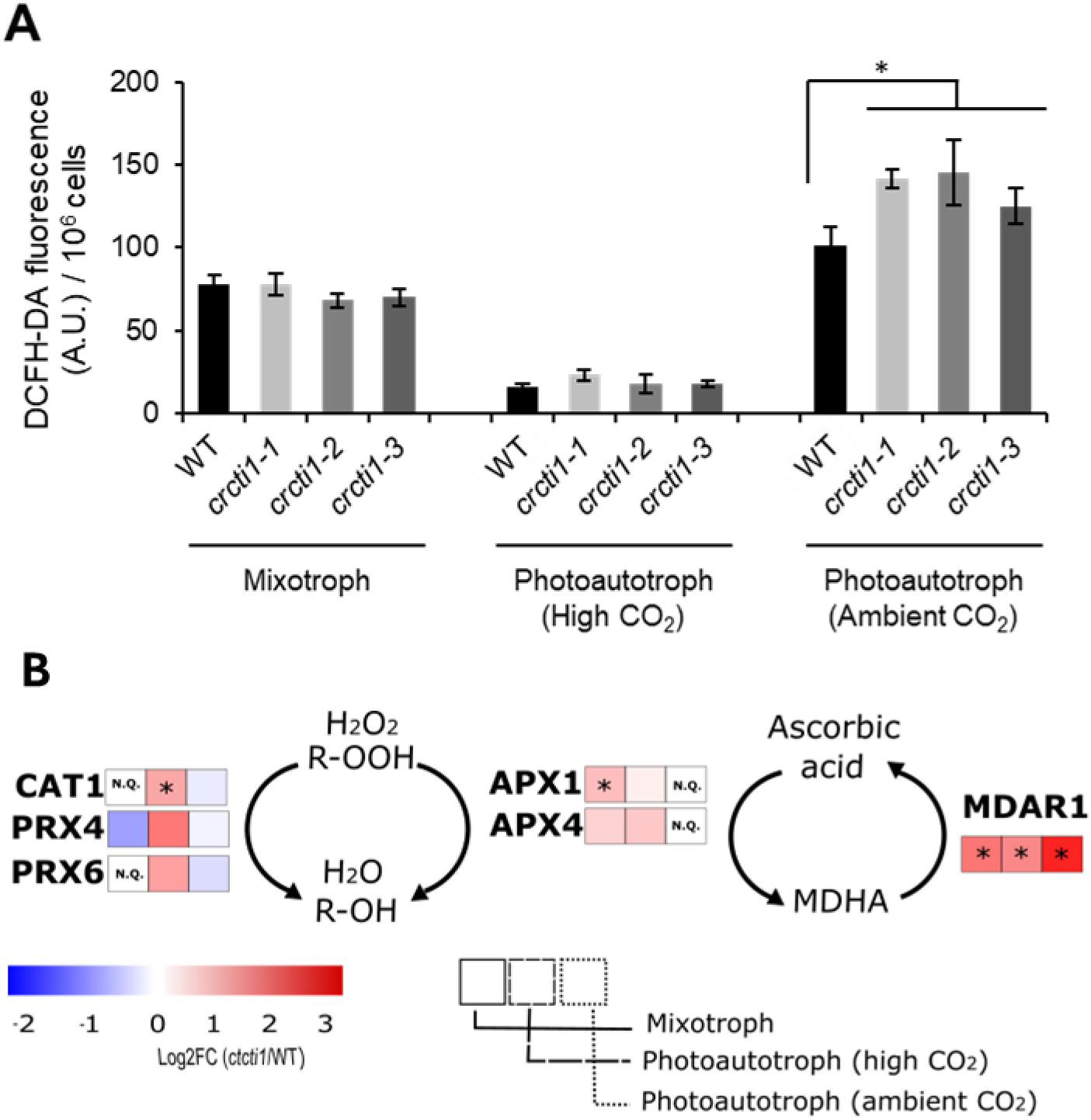
The highly abundant proteins in the enzymatic antioxidant defense pathway can effectively compensate for oxidative stress. **(A)** Intracellular ROS analysis by DCFH-DA staining of WT and *crcti1* mutants in mixotrophic and photoautotrophic condition (under air or supplemented with 2% CO_2_). All experiments were performed in three biological replicates (± S.D). * *p* < 0.05 (Student’s *t*-test). **(B)** Proteins detected by proteomics analysis. Fold-changes of protein amounts between *crcti1* mutants and wild-type were calculated as log2(*crcti1*/WT) and the significant differentially expressed proteins were displayed in the heat map. N.Q. represents protein not differently abundance with a p value ≤ 0.05. * represent two or more mutant lines with p-value ≤ 0.01, and similar fold-change values. Abbreviations: CAT1, catalase; PRX, peroxiredoxin; APX, ascorbate peroxidase; MDAR1, monodehydroascorbate reductase; MDHA, monodehydroascorbate; Asc, ascorbate.

Moreover, the chloroplast ascorbate peroxidases APX1 (Cre02.g087700) and APX4 (Cre05.g233900) were highly abundant in *crcti1* mutants under mixotrophic and photoautotrophic (high CO_2_) conditions. Especially, we observed that the other H_2_O_2_ scavengers peroxiredoxins PRX4 (Cre02.g080900) and PRX6 (Cre10.g422300), and catalase (CAT1 (Cre09.g417150) were highly abundant in *crcti1* mutants under photoautotrophic (high CO_2_) conditions. These results suggest that CrCTI1 deficiency may cause intracellular oxidative stress under ambient air, provoking an increase in major enzymes of the antioxidant pathways.

## DISCUSSION

Fatty acids are key building blocks for membrane formation. Their rate of synthesis should therefore be tightly controlled to meet with the varying needs of growth and division, which depend in particular on the environmental conditions plants and algae have to cope with (Thelen and Ohlrogge, 2002). This becomes even more important considering that i) free fatty acids are toxic and that its production should be coordinated to downstream metabolic needs (Wu *et al*., 2006); ii) plants and algae live in a challenging environment where light, CO_2_ and nutrient status may fluctuate rapidly. In this study, we identified and characterized the single plant CTI homolog i.e. CrCTI1 in the green microalga Chlamydomonas. We demonstrate that CrCTI1 acts as a negative regulator of de novo fatty acid synthesis in Chlamydomonas and that knocking out CrCTI1 in Chlamydomonas resulted in higher total lipid content in the cells regardless of N status but showed differential effects under different CO_2_ conditions. Here, we discuss the role of CrCTI1 under different carbon availability as well as its potential in terms of industrial applications.

### CrCTI1 regulates fatty acid synthesis in response to carbon availability

In Arabidopsis, it was reported that the negative regulation of fatty acid synthesis by CTI in the leaves is light dependent (Ye *et al*., 2020). As a photoautotrophic organism, Arabidopsis fix CO_2_ in the air as the sole carbon source for cell growth in the light, and thus light and ensuing carbohydrate production can be difficult to uncouple for diagnostic purposes. However, Chlamydomonas can easily be grown under photoautotrophic, mixotrophic, and even heterotrophic conditions (Harris, 2001); the cells can directly utilize external organic carbon like acetate as a carbon source in the dark without photosynthesis. Interestingly, a high TAG–accumulation phenotype was observed even in the dark for *crcti1* mutants, suggesting that CrCTI1 also functions in the absence of light, but presence of acetate. This suggests that the direct signal for CTI regulation could be carbon availability. We therefore propose that CrCTI1 is important in controlling fatty acid synthesis in both light and dark (acetate replete) conditions in microalgae, and this regulation is likely occurring through sensing carbon availability. Indeed, Wang et al (Wang *et al*., 2023) previously has observed that CrCTI1 – then named CPP1 – is present in a puncta that includes not only the Crα-CT (ACX1, Cre12.g519100) and Crβ-CT (BCX1, Cre12.g484000) proteins but also the chloroplastic isocitrate dehydrogenase 3 (IDH3) (Cre04.g214500). The authors have hypothesized a role of CrCTI1 in regulating acetyl-CoA metabolism downstream of fatty acid synthesis. It is worth mentioning, among many other potential interactors of CrCTI1 as pulled down by co-immunoprecipitation, one finds the CAS1 (Cre12.g497300), a known calcium (Ca^2+^)-binding protein playing a key role in regulating CO_2_ Concentrating Mechanism (CCM) (Wang *et al*., 2016) through a retrograde signal from the pyrenoid in the chloroplast to the nucleus. Indeed, many proteins known responsive to CO_2_ levels showed significant changes in the *crcti1* mutants (**Supplemental Table S1**), but whether CrCTI1 plays a role in this retrograde signaling pathway remain to be investigated. It would be interesting as well to decipher whether this carbon sensing is more a general mechanism present as well in land plants? This can be achieved by investigating the role of plant CTIs in non-photosynthetic tissues of plants (for example, white seeds, roots, tubers etc).

### The absence of CrCTI1 impacts central carbon metabolism in Chlamydomonas

Despite a higher oil content observed in the *crcti1* mutants, the protein abundance of enzymes involved in fatty acid synthesis and TAG accumulation was not significantly different between WT and the *crcti1* mutants. An increase in TAG accumulation due to hindering the glyoxylate cycle (Plancke *et al*., 2014) or β-oxidation (Kong *et al*., 2017, 2018) has been documented in several studies in Chlamydomonas. Isocitrate lyase null mutants in Chlamydomonas resulted in reduced acetate assimilation, with increased fatty acid content under mixotrophic condition, together with decreased abundance for enzymes involved in glyoxylate cycle, β-oxidation, and gluconeogenesis (Plancke *et al*., 2014). Similarly, *crcti1* mutants presented a decrease in the enzymes malate synthase (MAS1), citrate synthase 2 (CSY2), phosphoenolpyruvate carboxykinase 1 (PEPCK1), and peroxisomal acetyl-CoA synthase (ACS3) as described for *icl* null mutant. This finding suggests the higher increase in TAG observed for *crcti1* is partially explained by a reduction in glyoxylate cycle and β-oxidation, thus limiting TAG mobilization. Also, reduced levels for peroxisomal ACS3, while chloroplast ACS2 is enhanced under mixotrophic condition, suggests acetate is likely being pushed towards chloroplast ACS2 and de novo fatty acid synthesis. Consequently, we propose that *crcti1* creates an increased demand for carbon to sustain increased fatty acids and TAG levels, and carbon is supplied by acetate in the mixotrophic growth condition. This exogenous acetate carbon supply for fatty acid synthesis precludes the requirement for TAG mobilization, which was observed in the absence of acetate under photoautotrophic conditions.

### CrCTI1 controls carbon flux towards fatty acid production under ambient CO_2_ conditions

Under photoautotrophic conditions, CO_2_ is the sole carbon source for cell growth. Our observations indicate that CrCTI1 deficiency does not significantly impact cell growth when carbon supply is sufficient (i.e. at 2% CO_2_). However, under conditions of carbon limitation, CrCTI1 deficiency results in growth defects and increased TAG accumulation. Notably, the PSII efficiency remains unchanged compared to wild type, suggesting that CO_2_ availability is the critical factor influencing these outcomes. Cell growth is closely correlated with CO_2_ supply (Stitt, 1991). Rapidly growing and dividing cells under sufficient CO_2_ conditions require increased fatty acid synthesis to meet the demands for membrane lipid production. This suggests that negative regulators of ACCase activity, such as CrCTI1, may be downregulated or deactivated in response to abundant CO_2_. On the other hand, when the CO_2_ supply is limited, cells should control carbon flux precisely to maintain viability. From this perspective, *crcti1* mutants may lose control of fatty acid synthesis and utilize carbon sources required for the TCA cycle or central carbon metabolism. Therefore, CrCTI1, as a negative regulator of fatty acid synthesis, may be more important for cells under ambient CO_2_ conditions than under high CO_2_ conditions, and the fact that gene expression level of CrCTI1 increases under very low CO_2_ conditions supports this notion (Zheng *et al*., 2014).

### Manipulation of CrCTI1 may have advantages for large-scale lipid production

Nitrogen starvation is the most effective and traditional method of inducing oil accumulation, but it is not preferred in mass cultivation because it causes a stop in cell growth and moreover, it requires the replacement of the medium to induce nitrogen starvation (Pancha *et al*., 2014). Additionally, direct regulation (e.g., overexpression or repression of specific genes) of fatty acid metabolic pathways often have resulted in limited success (Wase *et al*., 2018). Interestingly, knocking out CrCTI1 resulted in increased TAG production without compromising cell growth during mixotrophic and heterotrophic conditions. A two-step production procedure can therefore be imagined for commercial oil production: firstly, growth during mixotrophic growth to allow cell growth and TAG accumulation; followed by a second phase where simply light is turned off to boost further TAG production. Based on our results, if applied to a switchable regulatory system using dCas9 (Vanegas *et al*., 2017), it could have a significant advantage in large-scale lipid production. Further understanding of the regulatory mechanism of CrCTI1 and its role in carbon sensing may enable more precise regulation.

## MATERIALS AND METHODS

### Strains and growth conditions

*Chlamydomonas reinhardtii* strains were maintained on Tris-Acetate-Phosphate (TAP) agar plates at 25°C under continuous illumination (30 μmol photons m^-2^ s^-1^). Cells in liquid medium were cultured in an incubator (INFORS HT), 120 rpm shaking, at 25°C, with 80 μmol photons m^-2^ s^-1^ illumination. Cells were cultivated either mixotrophically (TAP) or photoautotrophically in MOPS-buffered minimal media (MM) with or without an addition of 2% CO_2_ in the air, as specified in the text. To perform nitrogen (N) starvation, cells were centrifuged, and cell pellet was resuspended into media without nitrogen (TAP-N). Cell concentration was measured using a Beckman Coulter Multisizer 4 (Multisizer^TM^ Coulter Counter, Beckman Coulter, USA).

### Isolation of genomic DNA, RNA, and reverse transcription PCR analysis

Genomic DNA or RNA were extracted from exponentially grown Chlamydomonas cells based on the protocol of (Nguyen *et al*., 2013). To remove potential genomic DNA contaminations, the total RNA extracts were incubated with TURBO^TM^ DNase (Life technologies). The first strand cDNA was made using the SuperScript VILO cDNA Synthesis Kit (ThermoFisher Scientific). cDNA PCR was performed by using TB Green Premix Ex Taq II (Takara, Japan) on a 480 LightCycler thermocycler (Roche, USA). The gene-specific primers used for cDNA PCR were designed using the SnapGene software, which were listed in **Supplemental Table S2**. *RACK1* (receptor for activated kinase C 1) gene was used as an endogenous reference gene.

### Phylogenetic analysis

The amino acid sequences of the CTI family proteins from *Arabidopsis thaliana* (At) and Chlamydomonas (Cr) were aligned by using the Clustal W program built in Geneious Prime software. The phylogenetic tree was generated by using the neighbor-joining method with 5000 bootstrap replications. Molecular distances within the aligned sequences were calculated according to the position correction model.

### Generation of knock-out mutants using CRSPR-Cas9 RNP method

To generate CrCTI1 knock-out mutants, CRISPR-Cas9 RNP method was performed according to (Kim *et al*., 2020) with modifications. The single guide RNA (sgRNA) sequence was designed by Cas-Designer (http://www.rgenome.net/cas-designer) and selected as 5′-TAGCCAGCAGGTCCGCCCTG -3′. 100 μg of purified Cas9 protein (Cas9 expression plasmid: pET-NLS-Cas9-6xHis (Plasmid #62934)) and 70 μg of sgRNA synthesized by using GeneArt™ Precision gRNA Synthesis Kit (ThermoFisher, MA, USA), were mixed to form the RNP complex. For mutants screening, 0.5 μg of hygromycin-resistance (HygR; aphVII) gene cassette was co-transformed with RNP complex. Chlamydomonas transformation was performed in the 4 mm gap electroporating cuvette by electroporation with the specific parameter (600 V, 50 μF, infinity resistance). After 12 h recovery from transformation, cells were plated on TAP medium containing hygromycin (25 μg mL^-1^) and 1.5% agar.

### Yeast two-hybrid analysis

The sequences coding for the C-terminal region of CrCTI1 (amino acid 68-130) containing the coiled-coil domain and the full length of Crα-CT were cloned into the pGBKT7 DNA-BD vector. The sequences coding for the full length of Crα-CT and Crβ-CT were cloned into the pGADT7-AD vector. Primer sequences were given in Supplemental Table S2. Then, the yeast strain Y2HGold was co-transformed with different combinations of pGBKT7 DNA-BD and pGADT7-AD constructs using a modified LiOAc method (Ito *et al*., 1983). The yeast transformants plated with same volume on synthetic dropout (SD) medium lacking leucine and tryptophan, and SD medium lacking leucine, tryptophan, histidine, and adenine, were incubated at 30°C for 3 d.

### Lipid extraction, quantification by TLC and fatty acid composition analysis by GC-MS

Total lipids were extracted from exponentially grown phase Chlamydomonas cells by using a modified methyl tert-butyl ether (MTBE) method (Légeret *et al*., 2016). TAG and membrane lipids were quantified following a TLC based densitometry method as described previously in (Siaut *et al*., 2011). Fatty acid composition was analyzed after being converted to their fatty acid methyl esters (FAMEs) form. The FAMEs were then dissolved in hexane, transferred to a GC vial, and analyzed by a GC-FID apparatus (Agilent 7890A GC and Agilent 5975C MS, Agilent technologies) using a polar TR-WAX column (30 m x 0.25 mm x 0.50 μm). The GC conditions include split ratio of 1:20, injector and flame ionization detector temperature 240°C; oven temperature program 50°C for 2 min, then increasing at 15°C min^-1^ to 150°C, and then increasing again at 6°C min^-1^ to 240°C and holding at this temperature for 4 min. The flow rate of the carrier gas (H_2_) was 1 mL min^-1^.

### Protein extraction and in-solution digestion

To perform the proteomic study, cell cultures of the three *crcti1* mutant strains and the wild-type strain were harvested by centrifugation (1,500 *g*) for 5 min at the exponential phase. Each *crcti1* mutant line has five biologically independent cultures to ensure data reliability. Protein was extracted by mixing of 2.5 mL of Tris pH 8.8 buffered phenol and 2.5 mL of extraction media (0.1 M Tris-HCl pH 8.8, 10 mM EDTA, 0.4% 2-mercaptoethanol, 0.9 M sucrose) for 30 min at 4°C with agitating. Then the phenol phase was removed, and the aqueous phase was extracted again with 2.5 mL + 2.5 mL of extraction media and phenol. Phenol-extracted proteins were precipitated by adding 5 volumes of 0.1 M ammonium acetate in 100% methanol. Precipitated proteins were collected after centrifugation (10 min, 16,000 *g*, 4°C). Protein pellets were washed twice with 0.1 M ammonium acetate in methanol, and three times with 80% ice-cold acetone. An aliquot was removed and centrifuged at 16,000 *g*, 4°C for 5 min followed by pellet resuspension in urea buffer (6 M urea, 2 M thiourea, 100 mM ammonium bicarbonate).

Before applying the samples for mass spectrometry analysis, 10 μg of each sample was aliquoted and normalized to the same concentration and volume (40 µL) in urea buffer (6 M urea, 2 M thiourea, 100 mM ammonium bicarbonate). Reduction step (10 mM DTT in 10 mM ammonium bicarbonate) was performed at 30°C for 30 min followed by alkylation step (40 mM iodoacetamide (IAA) in 10 mM ammonium bicarbonate) for 1 hour in the dark at room temperature. Protein samples were diluted to a final urea concentration of 0.8 M before tryptic digestion. Digestion was performed by adding one volume of trypsin to reach a protein/enzyme ratio of 50:1 at 37°C for 16 h. To achieve optimal digestion, a second addition of trypsin with incubating for 4 h was performed. After digestion, the tryptic peptides were dried under vacuum centrifugation. Finally, samples were loaded onto Evotips according to the manufacturer’s instructions.

### Mass spectrometry data acquisition (UHPLC-MS/MS)

The mass spectrometric data was collected on an ultra-high-performance liquid chromatography (UHPLC) system (EvoSep) that was used for reverse-phase liquid chromatography for a 44 min gradient at 250 nL min^-1^ flow. Peptides were separated through a C18 analytical column (PepSep C18 Bruker Daltonics, 15 cm x 150 µm, 1.5 µm particle size). The mass spectra acquisition was performed alongside the chromatographic separation. The peptides were then subjected to analysis by a Data Dependent Acquisition (DDA) method. This analysis was conducted using a TimsTOF Pro 2 (Bruker Daltonics).

The instrument was operated in positive-ion and data-dependent PASEF mode with a m/z range from 100 to 1700. PASEF and TIMS were set to "on" for PASEF and TIMS. Ten PASEF and one MS frames were collected per cycle of 1.17 sec. Target MS intensity was set at 10,000 with a minimum threshold of 2500 from 20 to 59 eV, and a charge-state-based rolling collision energy table was employed. The method of an active exclusion/reconsider precursor with release after 0.4 min was used. A second MS/MS spectra were obtained if the precursor had a four-time increase in signal strength in subsequent scans (within a mass width error of 0.015 m/z).

### Protein identification and data analysis for global proteomics

To obtain *de novo* sequencing from MS/MS spectra, the PEAKS Studio 10.0 (Bioinformatics Solutions, Waterloo) software was used. PEAKS *de novo* sequencing was performed with precursor and fragment error tolerance values of 20 ppm and 0.01 Da, respectively. Trypsin was used as a protease with allowance of 3 maximum missed cleavage. Oxidation of methionine and carbamidomethylation of cysteine were set as variable and fixed modifications, respectively. A maximum of four variable modifications per peptide was allowed. PEAKS DB, which is a database search module in PEAKS Studio, was used in the second step to identify PSMs from existing protein databases (Zhang *et al*., 2012). The target-decoy method known as "decoy fusion" that is included in PEAKS was utilized to estimate the 1% FDR of the PEAKS DB result for establishing a confidence threshold for peptide spectrum matches (PSMs). The *C. reinhardtii* (Phytozome v6.1genome) was used as the reference database.

Data processing was performed using R studio with the package DEP 1.27.0. Intensity values were log2-transformed, and the quantitative data profiles were screened for missing values, which identified proteins had to be present in at least 70% of samples for further analysis. Data is background corrected and normalized by variance stabilizing transformation (vsn). Significantly different protein groups were evaluated by pairwise comparison analyses with Student’s t-test with 0.05 probability level with FDR correction.

### ROS measurement

Total ROS produced by Chlamydomonas cells was quantified by spectrophotometer. Cells were centrifuged at 1,500 *g*, 20°C for 5 min, resuspended in an equal volume of fresh culture medium, and then incubated with 5 µM DCFH-DA (Thermo Fisher Scientific) for 1 h in the dark at 25°C. After dark incubation, the cells were washed twice with culture medium to remove excess dye (Tran *et al*., 2019). The DCFH-DA fluorescence was measured using a Microplate reader (TecanM250) with an excitation of 485 nm and an emission of 530 nm. We calculated the total ROS by subtracting the value of background fluorescence (i.e. without DCFH-DA).

### Lipid droplet (LD) imaging

To observe LDs in Chlamydomonas, cells were harvested by centrifugation (500 *g*, 3 min), then resuspended in a small volume of fresh media. BODIPY (to a final concentration of 0.25 µg mL^-1^, from a stock of 50 µg mL^-1^ in DMSO) was then added to cell suspension and incubated in the dark for 15 min. Cells were then excited with a laser line at 488 nm, and emission was collected between 500-545 nm for BODIPY and between 650-730 nm for chlorophyll autofluorescence. All cell imaging were carried out under a 63x oil immersion objective in a confocal laser scanning microscope (LSM 780, Carl-ZEISS, Germany). Pseudo colors were applied using the ZEN (Carl-ZEISS) software.

## DATA AVAILABILITY STATEMENT

The mass spectrometry proteomics data have been deposited to the ProteomeXchange Consortium via the PRIDE (Perez-Riverol *et al*., 2022) partner repository with the dataset identifier PXD053580.

## ACKNOWLEDGEMENTS

We thank Stephan Cuine for excellent technical help. We also thank Dr. Jinsheng Zhu and Dr. Florian Veillet for helpful discussions on yeast two-hybrid analysis or on molecular biology techniques. We also thank the Zoom imaging facility. The funding of the HelioBiotec platform by the European Union Regional Developing Fund (ERDF), the Région Provence Alpes Côte d’Azur, the French Ministry of Research, and the CEA is also acknowledged.

## DISPLAY ELEMENTS

Figures 1 – 7

Supplemental files include:

Supplemental Figures S1-S6

Supplemental Table S1

Supplemental Table S2

## REFERENCE

Agrawal, G.K. and Thelen, J.J. (2006) Large Scale Identification and Quantitative Profiling of Phosphoproteins Expressed during Seed Filling in Oilseed Rape* S. Molecular & Cellular Proteomics 5, 2044–2059.

Bai, F., Yu, L., Shi, J., Li-Beisson, Y. and Liu, J. (2022) Long-chain acyl-CoA synthetases activate fatty acids for lipid synthesis, remodeling and energy production in Chlamydomonas. New Phytologist 233, 823–837.

Blom, N., Gammeltoft, S. and Brunak, S. (1999) Sequence and structure-based prediction of eukaryotic protein phosphorylation sites. Journal of molecular biology 294, 1351–1362.

Chapman, K.D. and Ohlrogge, J.B. (2012) Compartmentation of triacylglycerol accumulation in plants. Journal of Biological Chemistry 287, 2288–2294.

Chen, G. and Pramanik, B.N. (2009) Application of LC/MS to proteomics studies: current status and future prospects. Drug discovery today 14, 465–471.

Davis, M.S. and Cronan Jr, J.E. (2001) Inhibition of Escherichia coli acetyl coenzyme A carboxylase by acyl-acyl carrier protein. Journal of bacteriology 183, 1499–1503.

Eltayeb, A.E., Kawano, N., Badawi, G.H., Kaminaka, H., Sanekata, T., Shibahara, T., Inanaga, S. and Tanaka, K. (2007) Overexpression of monodehydroascorbate reductase in transgenic tobacco confers enhanced tolerance to ozone, salt and polyethylene glycol stresses. Planta 225, 1255–1264.

Gerhardt, E.C., Rodrigues, T.E., Müller-Santos, M., Pedrosa, F.O., Souza, E.M., Forchhammer, K. and Huergo, L.F. (2015) The bacterial signal transduction protein GlnB regulates the committed step in fatty acid biosynthesis by acting as a dissociable regulatory subunit of acetyl-CoA carboxylase. Molecular microbiology 95, 1025–1035.

Guchhait, R.B., Polakis, S.E., Dimroth, P., Stoll, E., Moss, J. and Lane, M.D. (1974) Acetyl coenzyme A carboxylase system of Escherichia coli: purification and properties of the biotin carboxylase, carboxyltransferase, and carboxyl carrier protein components. Journal of Biological Chemistry 249, 6633–6645.

Hajduch, M., Agrawal, G.K. and Thelen, J.J. (2007) Proteomics of seed development in oilseed crops. Plant Proteomics, 137–154.

Harris, E.H. (2001) Chlamydomonas as a model organism. Annual review of plant biology 52, 363–406.

Hu, Q., Sommerfeld, M., Jarvis, E., Ghirardi, M., Posewitz, M., Seibert, M. and Darzins, A. (2008) Microalgal triacylglycerols as feedstocks for biofuel production: perspectives and advances. The plant journal 54, 621–639.

Ito, H., Fukuda, Y., Murata, K. and Kimura, A. (1983) Transformation of intact yeast cells treated with alkali cations. Journal of bacteriology 153, 163–168.

Jeong, J., Baek, K., Yu, J., Kirst, H., Betterle, N., Shin, W., Bae, S., Melis, A. and Jin, E. (2018) Deletion of the chloroplast LTD protein impedes LHCI import and PSI–LHCI assembly in Chlamydomonas reinhardtii. Journal of experimental botany 69, 1147–1158.

Kim, J., Lee, S., Baek, K. and Jin, E. (2020) Site-specific gene knock-out and on-site heterologous gene overexpression in Chlamydomonas reinhardtii via a CRISPR-Cas9-mediated knock-in method. Frontiers in plant science 11, 306.

Kleinig, H. and Liedvogel, B. (1979) On the energy requirements of fatty acid synthesis in spinach chloroplasts in the light and in the dark. FEBS letters 101, 339–342.

Kong, F., Burlacot, A., Liang, Y., Légeret, B., Alseekh, S., Brotman, Y., Fernie, A.R., Krieger-Liszkay, A., Beisson, F. and Peltier, G. (2018) Interorganelle communication: peroxisomal MALATE DEHYDROGENASE2 connects lipid catabolism to photosynthesis through redox coupling in Chlamydomonas. The Plant cell 30, 1824–1847.

Kong, F., Liang, Y., Légeret, B., Beyly-Adriano, A., Blangy, S., Haslam, R.P., Napier, J.A., Beisson, F., Peltier, G. and Li-Beisson, Y. (2017) Chlamydomonas carries out fatty acid β-oxidation in ancestral peroxisomes using a bona fide acyl-CoA oxidase. The Plant Journal 90, 358–371.

Konishi, T. and Sasaki, Y. (1994) Compartmentalization of two forms of acetyl-CoA carboxylase in plants and the origin of their tolerance toward herbicides. Proceedings of the National Academy of Sciences 91, 3598–3601.

Lauersen, K.J., Willamme, R., Coosemans, N., Joris, M., Kruse, O. and Remacle, C. (2016) Peroxisomal microbodies are at the crossroads of acetate assimilation in the green microalga Chlamydomonas reinhardtii. Algal Research 16, 266–274.

Légeret, B., Schulz-Raffelt, M., Nguyen, H., Auroy, P., Beisson, F., Peltier, G., Blanc, G. and Li-Beisson, Y. (2016) Lipidomic and transcriptomic analyses of Chlamydomonas reinhardtii under heat stress unveil a direct route for the conversion of membrane lipids into storage lipids. Plant, cell & environment 39, 834–847.

Meyer, L.J., Gao, J., Xu, D. and Thelen, J.J. (2012) Phosphoproteomic analysis of seed maturation in Arabidopsis, rapeseed, and soybean. Plant physiology 159, 517–528.

Nguyen, H.M., Cuiné, S., Beyly-Adriano, A., Légeret, B., Billon, E., Auroy, P., Beisson, F., Peltier, G. and Li-Beisson, Y. (2013) The Green Microalga Chlamydomonas reinhardtii Has a Single ω-3 Fatty Acid Desaturase That Localizes to the Chloroplast and Impacts Both Plastidic and Extraplastidic Membrane Lipids. Plant Physiology 163, 914–928.

Nishi, H., Hashimoto, K. and Panchenko, A.R. (2011) Phosphorylation in protein-protein binding: effect on stability and function. Structure 19, 1807–1815.

Pancha, I., Chokshi, K., George, B., Ghosh, T., Paliwal, C., Maurya, R. and Mishra, S. (2014) Nitrogen stress triggered biochemical and morphological changes in the microalgae Scenedesmus sp. CCNM 1077. Bioresource technology 156, 146–154.

Perez-Riverol, Y., Bai, J., Bandla, C., García-Seisdedos, D., Hewapathirana, S., Kamatchinathan, S., Kundu, D.J., Prakash, A., Frericks-Zipper, A. and Eisenacher, M. (2022) The PRIDE database resources in 2022: a hub for mass spectrometry-based proteomics evidences. Nucleic acids research 50, D543–D552.

Plancke, C., Vigeolas, H., Höhner, R., Roberty, S., Emonds-Alt, B., Larosa, V., Willamme, R., Duby, F., Onga Dhali, D. and Thonart, P. (2014) Lack of isocitrate lyase in C hlamydomonas leads to changes in carbon metabolism and in the response to oxidative stress under mixotrophic growth. The Plant Journal 77, 404–417.

Roustan, V., Bakhtiari, S., Roustan, P.-J. and Weckwerth, W. (2017) Quantitative in vivo phosphoproteomics reveals reversible signaling processes during nitrogen starvation and recovery in the biofuel model organism Chlamydomonas reinhardtii. Biotechnology for biofuels 10, 1–24.

Ruiz-Sola, M.Á., Flori, S., Yuan, Y., Villain, G., Sanz-Luque, E., Redekop, P., Tokutsu, R., Küken, A., Tsichla, A. and Kepesidis, G. (2023) Light-independent regulation of algal photoprotection by CO2 availability. Nature communications 14, 1977.

Salie, M.J., Zhang, N., Lancikova, V., Xu, D. and Thelen, J.J. (2016) A Family of Negative Regulators Targets the Committed Step of de Novo Fatty Acid Biosynthesis. Plant Cell 28, 2312–2325.

Sauer, A. and Heise, K.-P. (1983) On the light dependence of fatty acid synthesis in spinach chloroplasts. Plant physiology 73, 11–15.

Shintani, D.K. and Ohlrogge, J.B. (1995) Feedback inhibition of fatty acid synthesis in tobacco suspension cells. The Plant Journal 7, 577–587.

Siaut, M., Cuiné, S., Cagnon, C., Fessler, B., Nguyen, M., Carrier, P., Beyly, A., Beisson, F., Triantaphylidès, C. and Li-Beisson, Y. (2011) Oil accumulation in the model green alga Chlamydomonas reinhardtii: characterization, variability between common laboratory strains and relationship with starch reserves. BMC biotechnology 11, 1–15.

Stitt, M. (1991) Rising CO2 levels and their potential significance for carbon flow in photosynthetic cells. Plant, Cell & Environment 14, 741–762.

Thelen, J.J. and Ohlrogge, J.B. (2002) Metabolic engineering of fatty acid biosynthesis in plants. Metabolic engineering 4, 12–21.

Tran, Q.G., Cho, K., Park, S.B., Kim, U., Lee, Y.J. and Kim, H.S. (2019) Impairment of starch biosynthesis results in elevated oxidative stress and autophagy activity in Chlamydomonas reinhardtii. Sci Rep 9, 9856.

Umezawa, T., Sugiyama, N., Takahashi, F., Anderson, J.C., Ishihama, Y., Peck, S.C. and Shinozaki, K. (2013) Genetics and phosphoproteomics reveal a protein phosphorylation network in the abscisic acid signaling pathway in Arabidopsis thaliana. Science signaling 6, rs8–rs8.

Vanegas, K.G., Lehka, B.J. and Mortensen, U.H. (2017) SWITCH: a dynamic CRISPR tool for genome engineering and metabolic pathway control for cell factory construction in Saccharomyces cerevisiae. Microbial cell factories 16, 1–12.

Wang, L., Patena, W., Van Baalen, K.A., Xie, Y., Singer, E.R., Gavrilenko, S., Warren-Williams, M., Han, L., Harrigan, H.R. and Hartz, L.D. (2023) A chloroplast protein atlas reveals punctate structures and spatial organization of biosynthetic pathways. Cell 186, 3499–3518.e3414.

Wang, L., Yamano, T., Takane, S., Niikawa, Y., Toyokawa, C., Ozawa, S.-i., Tokutsu, R., Takahashi, Y., Minagawa, J. and Kanesaki, Y. (2016) Chloroplast-mediated regulation of CO2-concentrating mechanism by Ca2+-binding protein CAS in the green alga Chlamydomonas reinhardtii. Proceedings of the National Academy of Sciences 113, 12586–12591.

Wase, N., Black, P. and DiRusso, C. (2018) Innovations in improving lipid production: Algal chemical genetics. Progress in lipid research 71, 101–123.

Williams, P.J.l.B. and Laurens, L.M. (2010) Microalgae as biodiesel & biomass feedstocks: Review & analysis of the biochemistry, energetics & economics. Energy & environmental science 3, 554–590.

Wu, J.-T., Chiang, Y.-R., Huang, W.-Y. and Jane, W.-N. (2006) Cytotoxic effects of free fatty acids on phytoplankton algae and cyanobacteria. Aquatic Toxicology 80, 338–345.

Ye, Y., Nikovics, K., To, A., Lepiniec, L., Fedosejevs, E.T., Van Doren, S.R., Baud, S. and Thelen, J.J. (2020) Docking of acetyl-CoA carboxylase to the plastid envelope membrane attenuates fatty acid production in plants. Nat Commun 11, 6191.

Yoshida, K., Igarashi, E., Mukai, M., Hirata, K. and Miyamoto, K. (2003) Induction of tolerance to oxidative stress in the green alga, Chlamydomonas reinhardtii, by abscisic acid. *Plant*, Cell & Environment 26, 451–457.

Young, D.Y. and Shachar-Hill, Y. (2021) Large fluxes of fatty acids from membranes to triacylglycerol and back during N-deprivation and recovery in Chlamydomonas. Plant Physiology 185, 796–814.

Zalutskaya, Z., Kharatyan, N., Forchhammer, K. and Ermilova, E. (2015) Reduction of PII signaling protein enhances lipid body production in Chlamydomonas reinhardtii. Plant Science 240, 1–9.

Zhang, J., Xin, L., Shan, B., Chen, W., Xie, M., Yuen, D., Zhang, W., Zhang, Z., Lajoie, G.A. and Ma, B. (2012) PEAKS DB: de novo sequencing assisted database search for sensitive and accurate peptide identification. Molecular & cellular proteomics 11.

Zheng, H.-Q., Chiang-Hsieh, Y.-F., Chien, C.-H., Hsu, B.-K.J., Liu, T.-L., Chen, C.-N.N. and Chang, W.-C. (2014) AlgaePath: comprehensive analysis of metabolic pathways using transcript abundance data from next-generation sequencing in green algae. BMC genomics 15, 1–12.

