## Supplemental figures and table S2 for "Knocking out the carboxyltransferase interactor 1 (CTI1) in Chlamydomonas boosted oil content by fivefold without affecting cell growth"

This file contains:

**Supplemental Figure S1 – S6**

**Supplemental Tables**

**Supplemental Table S1:** Proteomics data (see excel)

**Supplemental Table S2:** All primers used in this study

**Figure S1. Protein-protein interaction study on phosphorylation sites of CrCTI1 and Crα-CT.**

**(A)** Protein domain structures of *C. reinhardtii* CTI1 (CrCTI1) and α-CT (Crα-CT). The location of putative phosphorylation sites in Crα-CT was highlighted in red color. For the protein-protein interaction study, the coiled-coil domain of CrCTI1 (CrCTI1_CC) was used.

**(B)** Yeast two-hybrid (Y2H) assay.

**(C)** Yeast growth curve.

**(D)** Number of colony formed on SD/-Leu/-Trp and SD/-Leu/-Trp/-His/-Ade media. All putative phosphorylation sites in both CrCTI1 and Crα-CT (shown in (A) are mutated either to A or to D.

Abbreviations: AD, pGADT7 (prey vector); BD, pGBKT7 (bait vector); SD, synthetic drop-out medium supplements. T, threonine; Y, tyrosine; S, serine; D, aspartic acid; A, alanine. The yeast growth curve experiment was performed at least in three biological replicates (± S.D).

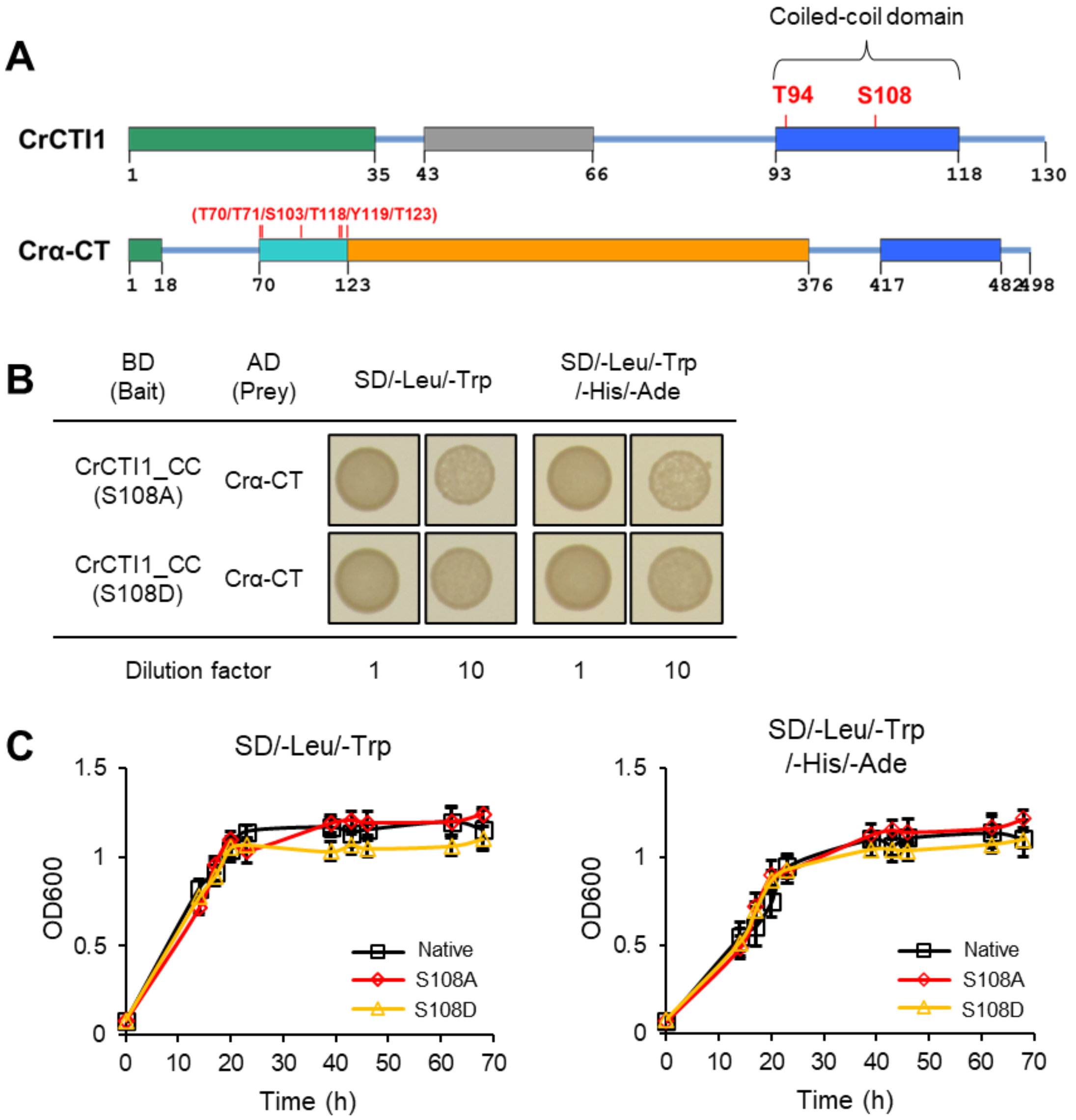


**D**

| BD (Bait) | AD (Prey) | Number of colony (per µg DNA) SD/-Leu/-Trp | Number of colony (per µg DNA) SD/-Leu/-Trp/-His/-Ade |
|---|---|---|---|
| CrCTI1_CC (Native) | Crα-CT_CC (Native) | 188 | 114 |
| CrCTI1_CC (Native) | Crα-CT (A) | 264 | 124 |
| CrCTI1_CC (Native) | Crα-CT (D) | 224 | 2 |
| CrCTI1_CC (A) | Crα-CT_CC (Native) | 192 | 120 |
| CrCTI1_CC (A) | Crα-CT (A) | 200 | 168 |
| CrCTI1_CC (A) | Crα-CT (D) | 292 | 10 |
| CrCTI1_CC (D) | Crα-CT_CC (Native) | 252 | 120 |
| CrCTI1_CC (D) | Crα-CT (A) | 256 | 122 |
| CrCTI1_CC (D) | Crα-CT (D) | 326 | 20 |

0 326

Number of colony (per µg DNA)

**Figure S2. Lipid profile under mixotrophic condition.**

**(A)** Mole percentage of fatty acids composition under mixotrophic condition.

**(B)** Membrane polar lipids content measured by TLC under mixotrophic N-repletion.

**(C)** TAG content measured by thin layer chromatography (TLC) under mixotrophic N-depletion. All experiments were performed in three biological replicates (± S.D). * $p < 0.05$, *** $p < 0.001$ (Student's *t*-test).

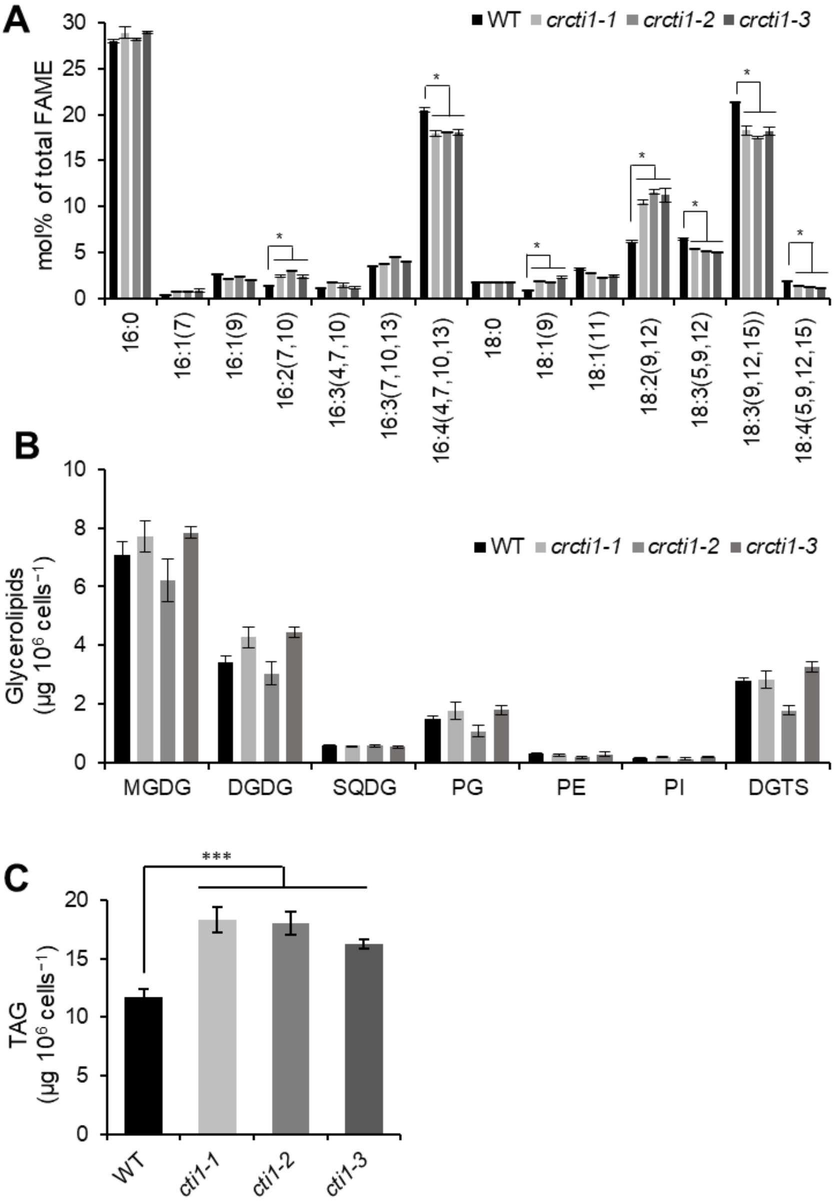

**Figure S3. Loss of CrCTI1 *in vivo* has no impact on photosynthesis under photoautotrophic conditions.**

**(A)** PSII yield measurement under photoautotrophic condition with 2% $CO_2$ supply. **(B)** PSII yield under photoautotrophic condition in air. All experiments were performed in three biological replicates (± S.D).

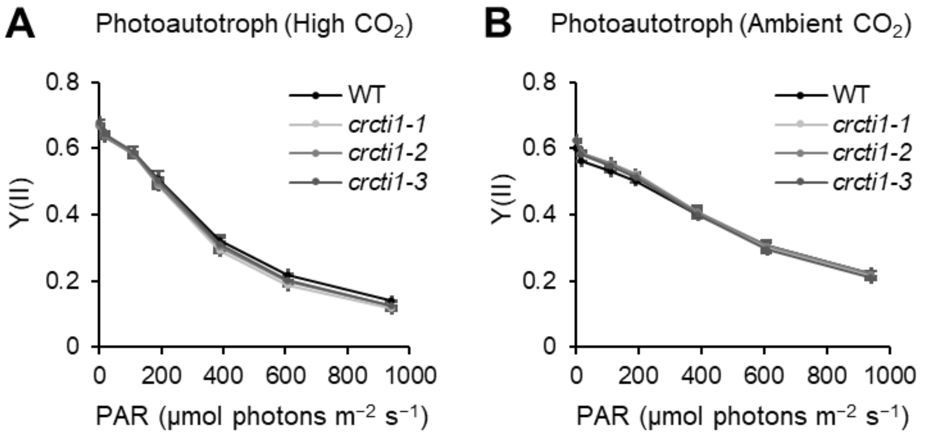

**Figure S4. Fatty acid composition.**

**(A)** Mole percentage of fatty acids composition under photoautotrophic condition with 2% $CO_2$ supply.

**(B)** Mole percentage of fatty acids composition under photoautotrophic condition in air. All experiments were performed in three biological replicates (± S.D). * $p < 0.05$ (Student's *t*-test).

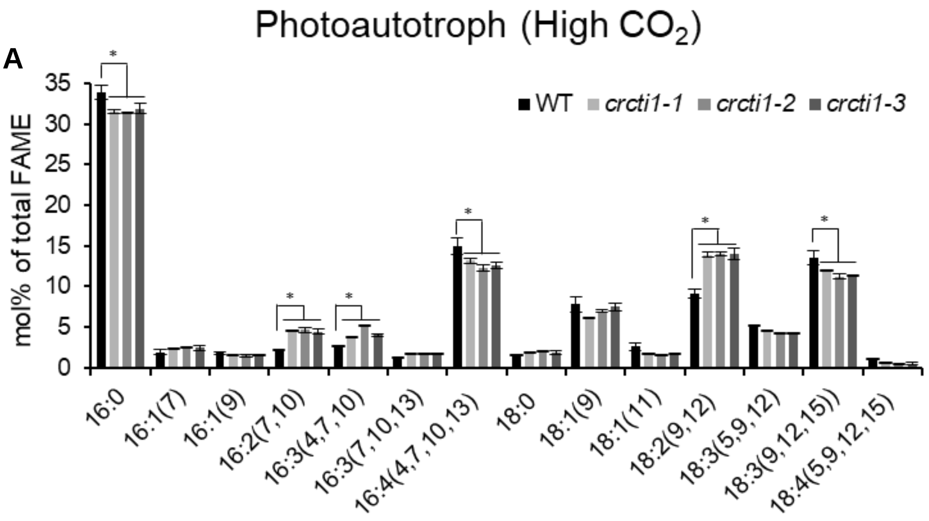


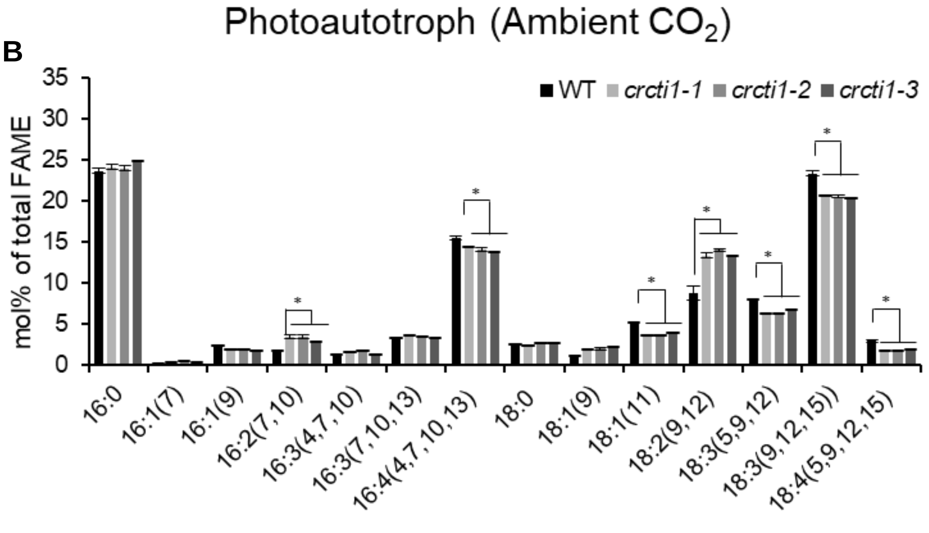

**Figure S5.**

Venn diagrams demonstrate the more (**A**) and less (**B**) abundant proteins in *crcti1* mutants. Only proteins differently expressed in a minimum of two mutant lines were, and with $p$ value ≤ 0.05 were included.

**A**

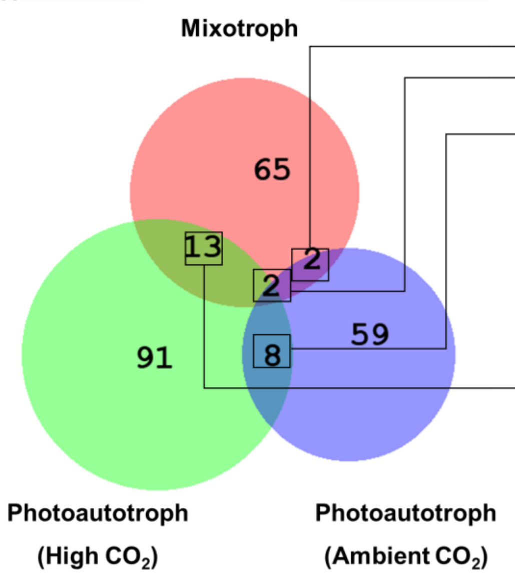


| ID | Gene | Arabidopsis ortholog | Protein description |
|---|---|---|---|
| Cre07.g356350 | DXS1 | | 1-deoxy-D-xylulose 5-phosphate synthase |
| Cre16.g674800 | NA | NA | NA |
| Cre17.g712100 | MDAR1 | AT1G53310 | Monodehydroascorbate reductase 1, peroxisomal |
| Cre12.g551950 | LTD1 | At1g50900 | Protein LHCP TRANSLOCATION DEFECT |
| Cre03.g144807 | MAS1 | AT5G03860 | Malate synthase |
| Cre10.g423250 | MDH2 | AT2G22780 | Malate dehydrogenase |
| Cre06.g251400 | MME6 | AT1G79750 | NADP-dependent malic enzyme |
| Cre04.g224750 | VAMP1 | AT3G54300 | Expressed Protein, similar to VAMP7 R-SNAREs |
| Cre07.g333950 | VPS45 | AT1G77140 | SM (Sec1-family) protein |
| Cre01.g004450 | CPLD42 | AT1G54520 | conserved expressed membrane protein |
| Cre12.g546050 | DXR1 | AT5G62790 | 1-deoxy-D-xylulose 5-phosphate reductoisomerase |
| Cre06.g295400 | BOU | | Mitochondrial substrate carrier protein |
| Cre03.g199535 | NA | AT4G17600 | Stress-enhanced protein 3 |
| Cre17.g726850 | NA | NA | Similar to Sjogren Syndrome Antigen A2 |
| Cre15.g643550 | PDX1 | AT5G01410 | Pyridoxin biosynthesis protein |
| Cre14.g623000 | NA | AT4G39970 | Phosphoglycolate phosphatase/2-deoxygucose-6-phosphate phosphatase |
| Cre13.g586450 | NA | NA | N-acetyltransferase |
| Cre06.g306300 | CHLI1 | AT4G18480 | Magnesium chelatase subunit I, isoform 1 |
| Cre05.g242000 | CHLD1 | AT1G08520 | Magnesium chelatase subunit D, chloroplast precursor |
| Cre01.g050950 | CHLP1 | AT1G74470 | Geranylgeranyl reductase CHLP |
| Cre02.g091050 | ALAD1 | AT1G69740 | Delta-aminolevulinic acid dehydratase |
| Cre07.g333150 | NA | NA | NA |
| Cre12.g497850 | NA | NA | NA |
| Cre13.g603850 | NA | NA | NA |
| Cre07.g352800 | NA | AT1G07910 | NA |

**More abundant in *cti1*/WT**

**B**

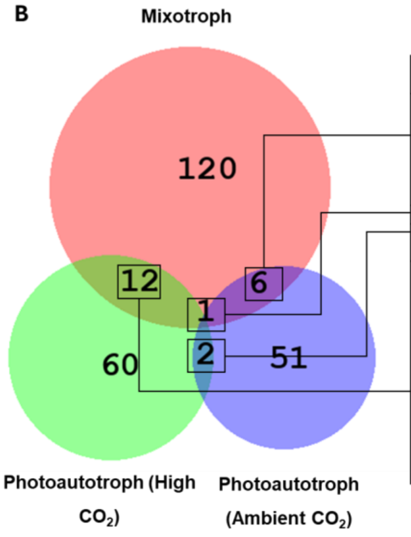


| ID | Gene | Arabidopsis ortholog | Protein description |
|---|---|---|---|
| Cre04.g217921 | COPB1 | AT4G31480 | Beta subunit of COP-I complex |
| Cre02.g081050 | FAP24 | AT1G65900 | Flagellar Associated Protein 24 |
| Cre14.g633750 | IPB2 | AT5G19820 | Importin beta-3 homolog |
| Cre11.g476650 | PUL1 | AT5G04360 | Pullulanase-type starch debranching enzyme 1 |
| Cre03.g160500 | TSK1 | AT3G11710 | Lysyl-tRNA synthetase |
| Cre06.g288700 | NA | AT4G36400 | NA |
| Cre16.g673852 | PPC1 | AT1G53310 | Phosphoenolpyruvate carboxylase 1 |
| Cre03.g172850 | OASTL3 | AT2G43750 | O-acetylserine (Thiol)-lyase/cysteine synthase 3 |
| Cre03.g145647 | | NA | NA |
| Cre02.g142800 | TRXZ1 | AT3G06730 | Thioredoxin z, chloroplastic |
| Cre12.g508750 | LHCA2 | AT1G19150 | Light-harvesting protein of photosystem I |
| Cre06.g274850 | HTR4 | NA | Histone H3 |
| Cre17.g714650 | HTR33 | AT4G40030 | Histone H3 |
| Cre03.g189950 | HOP1 | AT1G62740 | HSP70-HSP90 organizing protein |
| Cre17.g708800 | GSH2 | AT5G27380 | Glutathione synthetase |
| Cre16.g665364 | GDAK1 | AT1G12580 | Gliding Associated Kinase 1 |
| Cre09.g392867 | FMG1B | NA | Flagella Membrane Glycoprotein 1B |
| Cre16.g685573 | FAP214 | NA | Flagellar Associated Protein 214 |
| Cre12.g538100 | NA | AT4G19880 | Glutathione S-transferase |
| Cre12.g555450 | NA | AT5G07370 | inositol-tetrakisphosphate 5-kinase |
| Cre12.g512300 | NA | AT1G55020 | NA |

**Less abundant in *cti1*/WT**

**Figure S6.** 50 Most variable protein IDs ($p$ value < 0.05) in all three *crcti1* mutants against WT in mixotrophic growth **(A),** photoautotrophic (high $CO_2$) growth **(B)**, and photoautotrophic (ambient $CO_2$) growth **(C)**.

Abbreviations: ACC1, alpha-carboxyltransferase (ACCase complex); ADH1, dual function alcohol/acetaldehyde dehydrogenase; AGG3, aggregation 3; AIH2, agmatine iminohydrolase; ALS1, acetolactate synthase, large subunit; AMA3, alpha-amylase 3; AMA3, alpha-amylase 3; ANR1, anaerobiosis-related protein 1; ARG9, acetylornithine aminotransferase; ATPVC1, vacuolar ATP synthase subunit C; ATS1, ATP-sulfurylase; BTA1, diacylglyceryl-N,N,N-trimethylhomoserine synthesis protein; CAH4, mitochondrial carbonic anhydrase, beta type; CAH5, mitochondrial carbonic anhydrase; CHLD1, magnesium chelatase subunit D, chloroplast precursor; CHLM1, Mg protoporphyrin IX S-adenosyl methionine O-methyl transferase; CIS2, citrate synthase, glyoxysomal/microbody form; CPLD20, conserved in the green lineage and diatoms; CPLD49, cytochrome b6f biogenesis protein; CTAP4, chloroplast translocon associated protein 4; DEG1C, deg protease; DPE1, disproportionating enzyme 1; DXR1, 1-deoxy-D-xylulose 5-phosphate reductoisomerase, chloroplast precursor; ECH3, enoyl-CoA hydratase 1; EGD1, GAL4 DNA-binding enhancer protein 1; EPYC1, essential pyrenoid component 1; EPYC1, essential pyrenoid component 1; FAB2, plastid acyl-ACP desaturase; FAP214, flagellar associated protein 214; FAP24, flagellar associated protein 24; FAP295, flagellar associated protein 295; FAP303, flagellar associated protein 303; FAP332, flagellar associated protein 332; FAP346, FAS domain flagellar associated protein 346; FAP353, Rab-like GTP-binding flagellar associated protein 353; FAP354, Rab-like GTP-binding flagellar associated protein 354; FAP380, flagellar associated membrane protein 380; FAP380, flagellar associated membrane protein 380; FER1, pre-apoferritin; FLVB1, thylakoid flavodiiron protein; FMG1B, flagella membrane glycoprotein 1B; GAL1, galactose kinase; GAPC1, glyceraldehyde 3-phosphate dehydrogenase, chloroplastic; GBSS1A, granule-bound starch synthase IA; GCSP1, glycine cleavage system, P protein; GLC2B, glucosidase IIb; GSA1, glutamate-1-semialdehyde aminotransferase; HAV2, histone H2A variant; HID1, 3-hydroxyisobutyrate dehydrogenase; HLA8, transcriptional co-activator/nuclease induced by high light; HPR1, hydroxyl pyruvate reductase; ICL1, isocitrate lyase; IFR1, putative 2'-hydroxyisoflavone reductase; IPA1, importin alpha; KAT1, 3-oxoacyl CoA thiolase/acetyl-CoA acyltransferase 1; LC8, dynein arm light chain 8; LCI19, gamma hydroxybutyrate dehydrogenase; LCI3, low-$CO_2$-inducible protein; LCIB1, low-$CO_2$-inducible protein B; LCIC1, low-$CO_2$ inducible protein C; LCS3, long-chain acyl-CoA synthetase; LEU1L, osopropylmalate dehydratase, large subunit; LHCBM8, light-harvesting chlorophyll a/b binding protein of LHCII; LHCSR1, stress-related chlorophyll a/b binding protein 1; MAS1, malate synthase; MDAR1, monodehydroascorbate reductase 1; MDH2, malate dehydrogenase 2; MME6, NADP-dependent malic enzyme 6; NOP56, nucleolar protein, component of C/D snoRNPs; NOP58, nucleolar protein, component of C/D snoRNPs; NRX2, nucleoredoxin 2; NTR3, NADPH dependent thioredoxin reductase 3; NTR3, NADPH dependent thioredoxin reductase 3; NUOB10, NADH:ubiquinone oxidoreductase 17.8 kDa subunit; OGD1, 2-oxoglutarate dehydrogenase, E1 subunit; PDX1, pyridoxin biosynthesis protein; PMA2, P-type ATPase/cation transporter, plasma membrane; POB22, proteome of basal body 22; POR1, light-dependent protochlorophyllide reductase; PRPL7, chloroplast ribosomal protein L7/L12; RPN12, 26S proteasome regulatory subunit; RRA2, reduced residual arabinose 2k (SEC31) COP-II coat subunit; SECA1, chloroplast-associated SecA protein; SGA1, serine glyoxylate aminotransferase; SHMT2, serine hydroxymethyltransferase; STIC2, chloroplast protein biogenesis factor; THI4, thiazole biosynthetic enzyme; THIC1, hydroxymethylpyrimidine phosphate synthase; TSF2, phenylalanyl-tRNA synthetase; TSPSP1, trehalose-6-phosphate synthase/phosphatase, class I; VLE1, sporangin (Vegetative Lytic Enzyme 1); (VPS45) SM/Sec1-family protein.

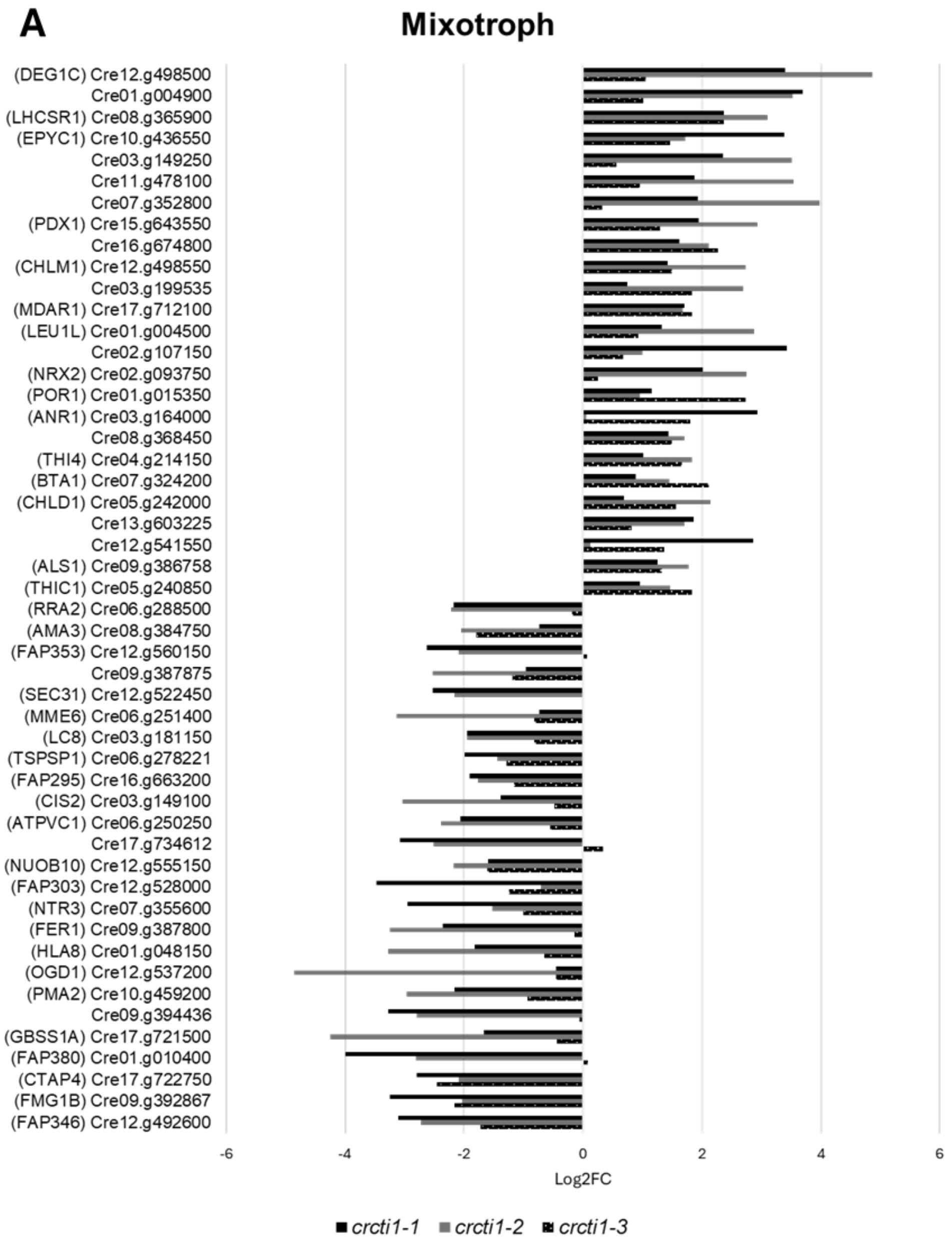
A
Mixotroph
(DEG1C) Cre12.g498500
Cre01.g004900
(LHCSR1) Cre08.g365900
(EPYC1) Cre10.g436550
Cre03.g149250
Cre11.g478100
Cre07.g352800
(PDX1) Cre15.g643550
Cre16.g674800
(CHLM1) Cre12.g498550
Cre03.g199535
(MDAR1) Cre17.g712100
(LEU1L) Cre01.g004500
Cre02.g107150
(NRX2) Cre02.g093750
(POR1) Cre01.g015350
(ANR1) Cre03.g164000
Cre08.g368450
(THI4) Cre04.g214150
(BTA1) Cre07.g324200
(CHLD1) Cre05.g242000
Cre13.g603225
Cre12.g541550
(ALS1) Cre09.g386758
(THIC1) Cre05.g240850
(RRA2) Cre06.g288500
(AMA3) Cre08.g384750
(FAP353) Cre12.g560150
Cre09.g387875
(SEC31) Cre12.g522450
(MME6) Cre06.g251400
(LC8) Cre03.g181150
(TSPSP1) Cre06.g278221
(FAP295) Cre16.g663200
(CIS2) Cre03.g149100
(ATPVC1) Cre06.g250250
Cre17.g734612
(NUOB10) Cre12.g555150
(FAP303) Cre12.g528000
(NTR3) Cre07.g355600
(FER1) Cre09.g387800
(HLA8) Cre01.g048150
(OGD1) Cre12.g537200
(PMA2) Cre10.g459200
Cre09.g394436
(GBSS1A) Cre17.g721500
(FAP380) Cre01.g010400
(CTAP4) Cre17.g722750
(FMG1B) Cre09.g392867
(FAP346) Cre12.g492600
-6
-4
-2
0
2
4
6
Log2FC
crcti1-1
crcti1-2
crcti1-3

**B**

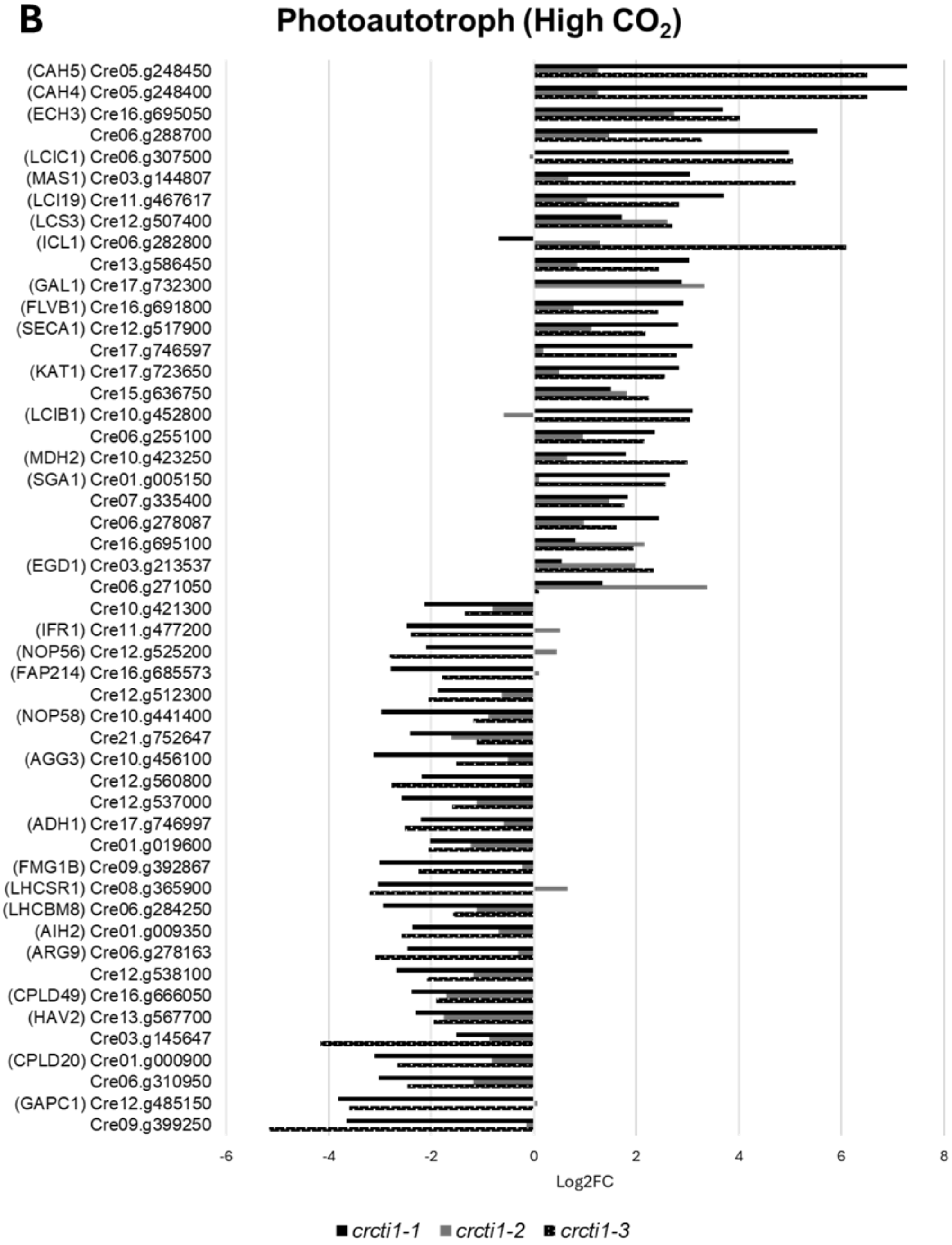

Photoautotroph (High $CO_2$)
(CAH5) Cre05.g248450
(CAH4) Cre05.g248400
(ECH3) Cre16.g695050
Cre06.g288700
(LCIC1) Cre06.g307500
(MAS1) Cre03.g144807
(LCI19) Cre11.g467617
(LCS3) Cre12.g507400
(ICL1) Cre06.g282800
Cre13.g586450
(GAL1) Cre17.g732300
(FLVB1) Cre16.g691800
(SECA1) Cre12.g517900
Cre17.g746597
(KAT1) Cre17.g723650
Cre15.g636750
(LCIB1) Cre10.g452800
Cre06.g255100
(MDH2) Cre10.g423250
(SGA1) Cre01.g005150
Cre07.g335400
Cre06.g278087
Cre16.g695100
(EGD1) Cre03.g213537
Cre06.g271050
Cre10.g421300
(IFR1) Cre11.g477200
(NOP56) Cre12.g525200
(FAP214) Cre16.g685573
Cre12.g512300
(NOP58) Cre10.g441400
Cre21.g752647
(AGG3) Cre10.g456100
Cre12.g560800
Cre12.g537000
(ADH1) Cre17.g746997
Cre01.g019600
(FMG1B) Cre09.g392867
(LHCSR1) Cre08.g365900
(LHCBM8) Cre06.g284250
(AIH2) Cre01.g009350
(ARG9) Cre06.g278163
Cre12.g538100
(CPLD49) Cre16.g666050
(HAV2) Cre13.g567700
Cre03.g145647
(CPLD20) Cre01.g000900
Cre06.g310950
(GAPC1) Cre12.g485150
Cre09.g399250
-6
-4
-2
0
2
4
6
8
Log2FC
crcti1-1
crcti1-2
crcti1-3


176

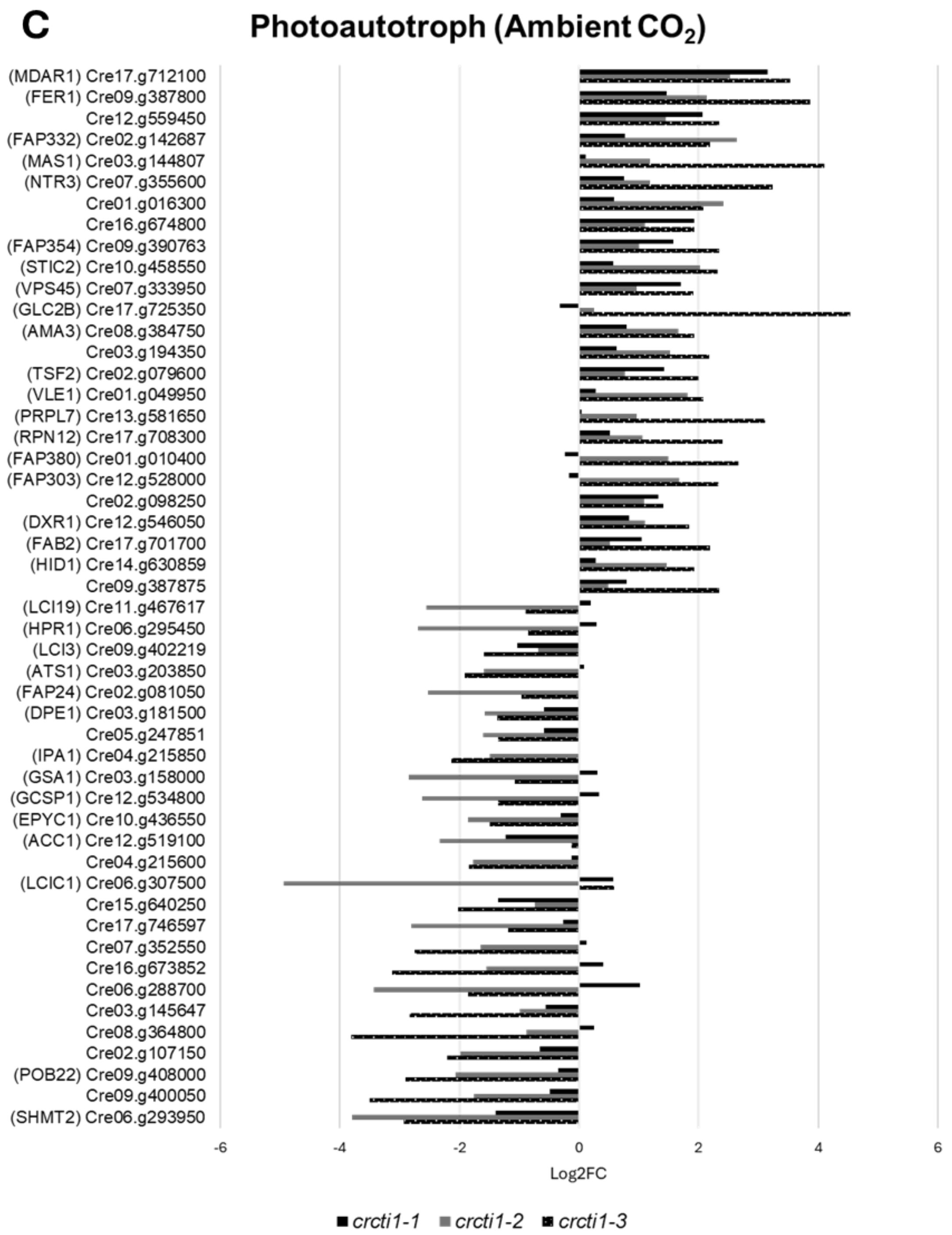
C
Photoautotroph (Ambient $CO_2$)
(MDAR1) Cre17.g712100
(FER1) Cre09.g387800
Cre12.g559450
(FAP332) Cre02.g142687
(MAS1) Cre03.g144807
(NTR3) Cre07.g355600
Cre01.g016300
Cre16.g674800
(FAP354) Cre09.g390763
(STIC2) Cre10.g458550
(VPS45) Cre07.g333950
(GLC2B) Cre17.g725350
(AMA3) Cre08.g384750
Cre03.g194350
(TSF2) Cre02.g079600
(VLE1) Cre01.g049950
(PRPL7) Cre13.g581650
(RPN12) Cre17.g708300
(FAP380) Cre01.g010400
(FAP303) Cre12.g528000
Cre02.g098250
(DXR1) Cre12.g546050
(FAB2) Cre17.g701700
(HID1) Cre14.g630859
Cre09.g387875
(LCI19) Cre11.g467617
(HPR1) Cre06.g295450
(LCI3) Cre09.g402219
(ATS1) Cre03.g203850
(FAP24) Cre02.g081050
(DPE1) Cre03.g181500
Cre05.g247851
(IPA1) Cre04.g215850
(GSA1) Cre03.g158000
(GCSP1) Cre12.g534800
(EPYC1) Cre10.g436550
(ACC1) Cre12.g519100
Cre04.g215600
(LCIC1) Cre06.g307500
Cre15.g640250
Cre17.g746597
Cre07.g352550
Cre16.g673852
Cre06.g288700
Cre03.g145647
Cre08.g364800
Cre02.g107150
(POB22) Cre09.g408000
Cre09.g400050
(SHMT2) Cre06.g293950
-6
-4
-2
0
2
4
6
Log2FC
crcti1-1
crcti1-2
crcti1-3

### Supplemental Tables

**Supplemental Table S1:** Proteomics data (see excel)

**Supplemental Table S2:** All primers used in this study

| | **Primer forward (5'-3')** | **Primer reverse (5'-3')** |
|---|---|---|
| **Gene cloning** | | |
| CrCTI1 | ATGTCTGCCATGCTCCGTCAAC | TTACGGCGTCGACTCAATCTTCAT |
| **Yeast two-hybrid with restriction sites** | | |
| CrCTI1 | CCGGAATTCAAGGCGCTCCTAGGC | CGCGGATCCCGGCGTCGA |
| Crα-CT | CCGGAATTCATGCAGGTCCTCAAGAGCA | CGCGGATCCAGCGGCGGC |
| Crβ-CT | CCGGAATTCATGGCGCCGCAG | CCGCTCGAGCTTCATGCGGAACA |
| T7-F | TAATACGACTCACTATAGGGC | |
| BD-R | | TTTTCGTTTTAAAACCTAAGAGTC |
| AD-R | | AGATGGTGCACGATGCACAG |
| **Site-directed mutagenesis** | | |
| CrCTI1 S108A | CGCCGCGATCGACGAGGTGGCCGCG | CGATCGCGGCGTTGAGCTGAGCAATCTTCT |
| CrCTI1 S108D | CGCCGACATCGACGAGGTGGCCGCG | CGATGTCGGCGTTGAGCTGAGCAATCTTCT |
| **CRISPR-Cas9** | | |
| CRISPR sgRNA | TAATACGACTCACTATAGGCTGGCTACGCGCAGCAGGC | TTCTAGCTCTAAAACGCCTGCTGCGCGTAGCCAGC |
| crcti1 screen | GGATCGATGCACATGAATTCTAGC | TATGTGTGCGATTCCACGCAAGC |
| **RT-PCR** | | |
| CrCTI1 | CCCTCAGATCTCCAAGGC | CGTTGAGCTGAGCAATCTTC |
| RACK1 | CATAATTTCGGCCTCCCGTGA | AGTGTCCGTAGAGACGCTTCT |
